# Investigating structural variant, indel and single nucleotide polymorphism differentiation between locally adapted Atlantic salmon populations

**DOI:** 10.1101/2023.09.12.557169

**Authors:** Laurie Lecomte, Mariann Árnyasi, Anne-Laure Ferchaud, Matthew Kent, Sigbjørn Lien, Kristina Stenløkk, Florent Sylvestre, Claire Mérot, Louis Bernatchez

**Affiliations:** Institut de Biologie Intégrative et des Systèmes (IBIS), Université Laval; Département de biologie, Université Laval; Department of Animal and Aquacultural Sciences (IHA), Faculty of Life Sciences (BIOVIT), Centre for Integrative Genetics (CIGENE), Norwegian University of Life Sciences (NMBU); Parks Canada, Office of the Chief Ecosystem Scientist, Protected Areas Establishment; Institut de Biologie Intégrative et des Systèmes (IBIS), Université Laval; Département de biologie, Université Laval; UMR 6553 Ecobio, OSUR, CNRS, Université de Rennes; Institut de Biologie Intégrative et des Systèmes (IBIS), Université Laval; Département de biologie, Université Laval

## Abstract

Genomic structural variants (SVs) are now recognized as an integral component of intraspecific polymorphism and are known to contribute to evolutionary processes in various organisms. However, they are inherently difficult to detect and genotype from readily available short-read sequencing data, and therefore remain poorly documented in wild populations. Salmonid species displaying strong interpopulation variability in both life history traits and habitat characteristics, such as Atlantic salmon (*Salmo salar*), offer a prime context for studying adaptive polymorphism, but the contribution of SVs to fine-scale local adaptation has yet to be explored. Here, we performed a comparative analysis of SVs, single nucleotide polymorphisms (SNPs) and small indels (< 50 bp) segregating in the Romaine and Puyjalon salmon, two putatively locally adapted populations inhabiting neighboring rivers (Québec, Canada) and showing pronounced variation in life history traits, namely growth, fecundity, and age at maturity and at smoltification. We first catalogued polymorphism using a hybrid SV characterization approach pairing both short (16X) and long-read sequencing (20X) for variant discovery with graph-based genotyping of SVs across 60 salmon genomes, along with characterization of SNPs and small indels from short reads. We thus identified 115,907 SVs, 8,777,832 SNPs and 1,089,321 short indels, with SVs covering 4.8 times more base pairs than SNPs. All three variant types revealed a highly congruent population structure and similar patterns of *F_ST_*and density variation along the genome. Finally, we performed outlier detection and redundancy analysis (RDA) to identify variants of interest in the putative local adaptation of Romaine and Puyjalon salmon. Genes located near these variants were enriched for biological processes related to nervous system function, suggesting that observed variation in traits such as age at smoltification could arise from differences in neural development. This study therefore demonstrates the feasibility of large-scale SV characterization and highlights its relevance for salmonid population genomics.

## 1 Introduction

Differences in DNA sequence and structure among individuals within species, referred to as genetic variation, serve as the basis for key evolutionary mechanisms such as speciation and local adaptation (Barrett & Schluter, 2008). Genetic variation can be described as a wide spectrum of variants of various sizes, ranging from single nucleotide polymorphisms (SNPs) to larger structural variants (SVs), which may span megabase-long stretches of DNA or even whole chromosomes (Feuk, Carson & Scherer, 2006; Wellenreuther & Bernatchez, 2018; Mérot et al., 2020). SVs such as insertions, deletions, duplications and inversions are now recognized as the main component of genetic variation, as they affect at least two to eight times more bases in genomes than SNPs (Catanach et al., 2019; Mérot et al., 2020; Hämälä et al., 2021). This estimate tends to increase as our ability to detect SVs from high-throughput sequencing data is constantly improving (Ho, Urban & Mills, 2020; Mérot et al., 2020).

SVs are also known to have a broad range of consequences at various biological levels. At the molecular scale, they may influence gene dosage, gene expression, DNA interactions and tridimensional structure by altering genetic elements’ proximity and copy number (Feuk et al., 2006; Gamazon & Stranger, 2015; Spielmann, Lupiáñez & Mundlos, 2018). SVs that disrupt collinearity between homologous chromosomes, especially large inversions, are also likely to restrict or suppress recombination (Crown et al., 2018; Rowan et al., 2019). This may result in an apparent reduced gene flow around SVs, which may link co-adapted alleles, thus promoting the formation of supergenes that may underlie complex and adaptive phenotypes (Rieseberg, 2001; Kirkpatrick & Barton, 2006; Thompson & Jiggins, 2014).

A growing body of evidence also suggests that SVs can be involved in evolutionary mechanisms in various species (reviewed in Wellenreuther & Bernatchez, 2018). For instance, supergenes arising from large inversions have been linked to adaptive variation in wing color patterns in *Heliconius* butterfly (Joron et al., 2011), to migratory behavior in rainbow trout (*Oncorhynchus mykiss*; Pearse et al., 2014; Pearse et al., 2019) and Atlantic cod (*Gadus morhua*; Kirubakaran et al., 2016; Berg et al., 2017), as well as to reproductive strategies in the ruff (*Philomachus pugnax*; Küpper et al., 2015) and the white-throated sparrow (*Zonotrichia albicollis*; Tuttle, 2003). Other key examples of inversion polymorphism involved in ecotype divergence and local adaptation have been documented in the seaweed fly (*Coelopa frigida*; Mérot et al., 2018) and in three-spined stickleback (*Gasterosteus aculeatus*; Jones et al., 2012). Besides large inversions, copy number variants (CNVs) have also been linked to adaptation to local temperature regimes in American lobster (*Homarus americanus*; Dorant et al., 2020) and to glacial lineage divergence in capelin (*Mallotus villosus*; Cayuela et al., 2021). Industrial melanism in peppered moths (*Biston betularia*), a textbook example of rapid adaptation to environmental change, has been associated with an intronic insertion in the *cortex* gene (Van’t Hof et al., 2016). Similarly, a 2.25-kb intronic insertion would explain color pattern divergence among lineages in the *Corvus* genus, promoting reproductive isolation and thus leading to speciation (Weissensteiner et al., 2020).

Despite such well-documented cases of adaptive genomic rearrangements, most SVs other than large inversions remain understudied in a population genomics context. Relative to SNPs, very little is known about how such a large component of genetic variation is distributed within and between wild populations. Indeed, while standard procedures and pipelines are available for population-scale SNP calling, SV detection and genotyping, on the other hand, involve significant challenges for large, multisample datasets (Mahmoud et al., 2019; Ho, Urban & Mills, 2020).

Calling SVs requires specialized software due to their complexity and diversity in type and length. SV callers rely on various signals of discordance in read mapping relative to a reference genome in order to infer SVs in a given sample (Lin et al., 2015; Mahmoud et al., 2019). Because short-read sequencing is widely available and affordable, it is an appropriate technology for population-scale study of genetic variation. However, the performance of short-read-based SV callers is highly variable (Cameron, Di Stefano & Papenfuss, 2019). They are known to lack sensitivity, as true positive detection rates can be as low as 10% (Huddleston et al., 2017; Sedlazeck et al., 2018a). They also show low precision, with high false discovery rates reaching 89% for some datasets (Mills et al., 2011), especially for calls near SNPs, indels, low-complexity regions and repeats (Cameron, Di Stefano & Papenfuss, 2019). In fact, short reads are hard to map to the reference genome owing to their small length, especially when they include numerous sequencing errors, repeats (Sedlazeck et al., 2018b) or sequences differing considerably from the reference, such as SVs. Spurious mapping may result in underreporting of variation, an issue known as reference allele bias (Nielsen et al., 2011; Brandt et al., 2015). Calling SVs in a given dataset using multiple callers (ensemble calling), may increase the range of SV types and sizes detected or reduce false discovery rate compared to single-tool SV calling. For instance, SV callsets may be merged across callers, then filtered for calls supported by a given minimum number of tools (Auton et al., 2015). However, the improvement in sensitivity and/or precision strongly depends on the callers used in combination (Kosugi et al., 2019; Mahmoud et al., 2019).

By contrast, recent advances in third generation sequencing platforms (Oxford Nanopore and Pacific Biosciences’ technologies) have brought significant improvements regarding SV calling. Long reads span kilobase-long segments of DNA, thus fully overlapping SVs and their breakpoints, which considerably facilitates read mapping (Sedlazeck et al., 2018b) and improves sensitivity of SV detection (Mahmoud et al., 2019), especially for novel insertions (Ho, Urban & Mills, 2020). Specialized algorithms and pipelines have been developed to process long reads and account for their length and higher sequencing error rate (Rang, Kloosterman & de Ridder, 2018; Delahaye et al., 2021). However, high costs prevent long-read sequencing from becoming a routine tool for population-scale SV studies for species with large genomes such as salmonid fishes (usually around 3 Gb), which requires the sequencing of many genomes, namely for accurate estimates of allele frequency.

To provide an adequate balance between accurate SV characterization and genotyping in large datasets, emerging hybrid approaches can be considered, such as pairing affordable short-read sequencing for all samples with high performance third generation sequencing for a small subset of genomes only. Candidate SVs called from long reads can then be genotyped in all samples from short-read data using pangenome graphs, which offer considerable advantages over conventional, linear reference-based methods. Indeed, in a reference pangenome, the reference genome is represented as a base graph structure where known variants and alternate alleles are encoded as alternate paths, i.e., series of nodes and links (Paten et al., 2017). The integration of known genetic variation within the reference greatly facilitates mapping of reads that overlap such variants, thus improving both SV detection and genotyping, and reducing reference allele bias (Ameur, 2019). This approach has shown promising results for genome-wide population-scale SV detection in human (*Homo sapiens*; Yan et al., 2021), soybean (*Glycine max*; Lemay et al., 2022), lake whitefish (*Coregonus clupeaformis*; Mérot et al., 2023) <u>and in kākāpō parrots (*Strigops habroptilus*; Wold et al., 2023)</u>.

Knowledge pertaining to SVs remains minimal in salmonid fishes, despite their genomes being extensively studied for aquaculture applications. The first comprehensive catalog of genome-wide SVs for Atlantic salmon (*Salmo salar*) was produced by Bertolotti et al. (2020) by calling putative SVs using short-read-based caller LUMPY (Layer et al., 2014) in 492 wild and domestic salmon from various populations in Europe and North America. SV calls were then manually curated with SV-plaudit (Belyeu et al., 2018) in order to eliminate false positives, yielding 15,483 high confidence SVs matching the expected population structure. This study also revealed a subset of outlier SVs overlapping genes enriched for brain expression, suggesting an implication in salmon domestication. Other population-scale SV catalogs were published for the rainbow trout (*Oncorhynchus mykiss*; Liu et al., 2021), and two sympatric sister species of lake whitefish (*Coregonus* sp.; Mérot et al., 2023).

Further work is required in order to fully appreciate SVs’ relevance in the genomics and biology of Atlantic salmon, which could serve as an ideal candidate species for studying adaptive SVs and developing an efficient population-scale SV discovery pipeline. SVs and larger chromosomal rearrangements are likely a key feature of salmonid genomes, as they are critical to the reploidization process following the salmonid-specific fourth vertebrate whole-genome duplication that occurred at least 60 million years ago (Ss4R) (Allendorf & Thorgaard, 1984; Crête-Lafrenière, Weir & Bernatchez, 2012; Lien et al., 2016). Sequence repeats, which account for 50 to 60% of the Atlantic salmon genome (de Boer et al., 2007), are also known to promote SV formation (Levy-Sakin et al., 2019). Moreover, Atlantic salmon display considerable life history trait variation both within and between wild populations (Klemetsen et al., 2003). Consequently, there is considerable interest in understanding the genetic architecture of traits such as growth rate and disease resistance for aquaculture (Gjedrem & Rye, 2018), but also in the context of local adaptation, which usually involves such life history trait variation (Taylor, 1991; Lu & Bernatchez, 1999; Fraser & Bernatchez, 2005). Local adaptation is expected to be a major driver of population structure in Atlantic salmon, given its homing behavior (Allendorf & Waples, 1996) and the variability in habitat conditions (Taylor, 1991; Kawecki & Ebert, 2004). Indeed, the association between the genetic structure of seven groups of local salmon populations in Eastern Canada and regional rivers’ environmental parameters suggests adaptive SNP divergence among these groups (Dionne et al., 2008; Bourret et al., 2013). Previous studies have also highlighted a few adaptive large chromosomal rearrangements in wild Atlantic salmon populations (Wellband et al., 2019; Watson et al., 2022), as well as divergent SVs between domestic and wild populations (see Bertolotti et al., 2020). However, SVs’ contribution to local adaptation remains poorly documented among North American populations, especially at a finer geographic scale (e.g., within neighboring rivers).

Two parapatric Atlantic salmon populations from the Romaine and Puyjalon rivers (Québec, Canada; 50.306337, -63.795602; Figure 1B) represent a prime case of putative fine-scale local adaptation. Indeed, admixture analysis and fixation index calculation (*F_ST_* = 0.036) based on microsatellite markers showed moderate differentiation between Romaine (RO) and Puyjalon (PU) salmon, despite their geographical proximity and habitat connectivity (Albert & Bernatchez, 2006). Furthermore, they exhibit different trade-offs in major life history traits: earlier age at smoltification and sexual maturity have been reported among wild Romaine salmon (Fontaine et al., 2000; Belles-Isles et al., 2004; WSP Global, 2019), as well as in wild-born Romaine salmon reared in a hatchery environment at the LARSA (Laboratoire de Recherche en Sciences Aquatiques; Université Laval, Québec), whereas wild-born Puyjalon salmon have shown higher growth rates over several cohorts in the same hatchery conditions (Therrien et al., 2017; Langlois-Parisé et al., 2018; T. Dion et al., 2020; T. Dion, Langlois-Parisé & Proulx, 2020). The persistence of such life history trait variation among cohorts in both wild and controlled environments strongly suggests heritable genetic variation likely linked to local adaptation, as the Romaine and Puyjalon rivers differ in spawning habitat quality, substrate and hydrological parameters (Schieffer, 1975; Fontaine et al., 2000; GENIVAR, 2002; Belles-Isles et al., 2004; WSP Global, 2019). However, the genetic basis of this putative local adaptation has yet to be investigated.

**Figure 1.**
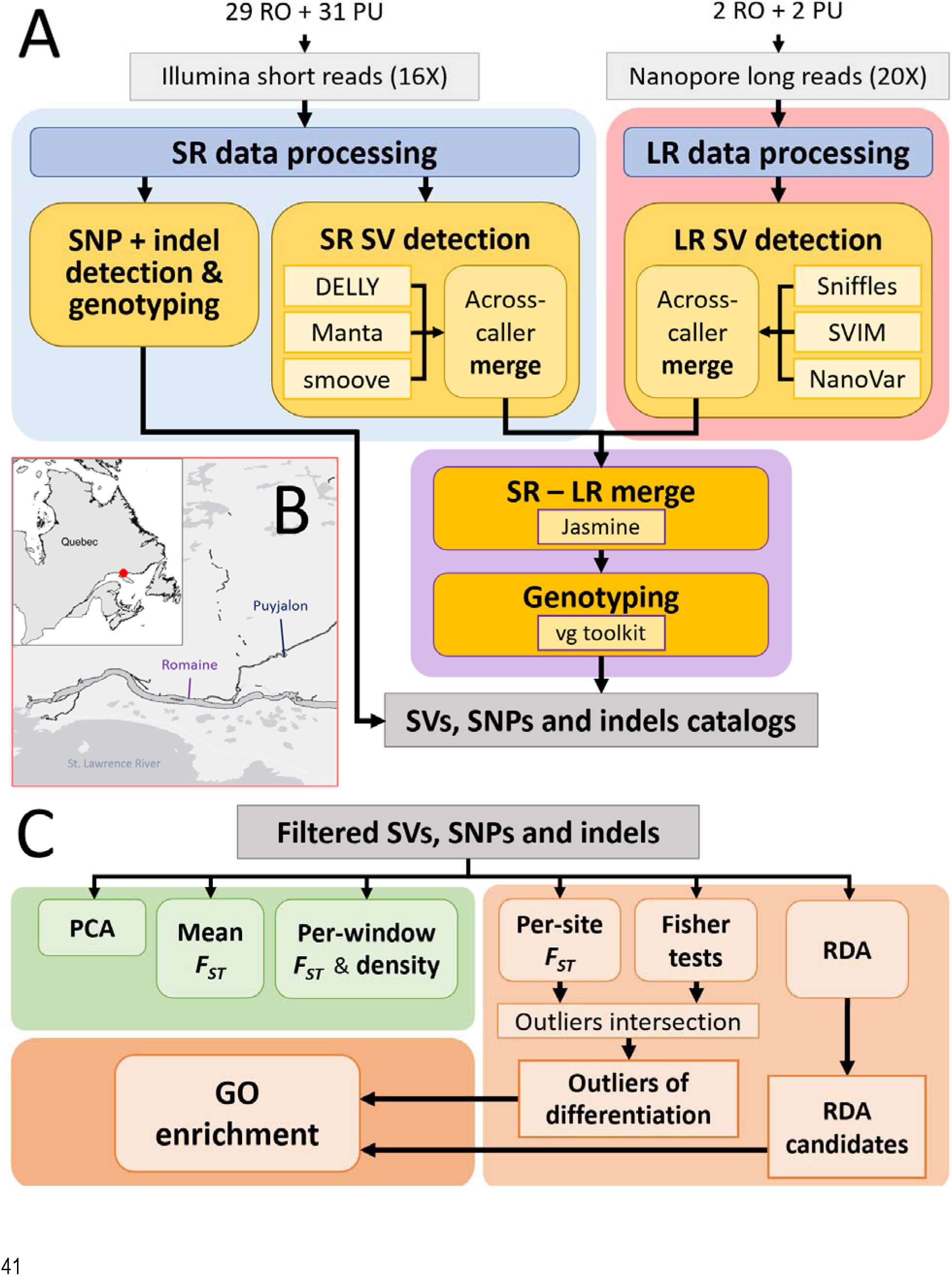
Overview of **A)** polymorphism detection pipelines used for population-scale characterization of structural variants (SVs), single nucleotide polymorphisms (SNPs) and small indels within the genomes of Romaine (RO) and Puyjalon (PU) salmon (SR: short reads; LR: long reads), **B)** location of the Romaine and Puyjalon rivers (red dot) in Québec, Canada, and **C)** comparative genomics analyses performed on catalogued variants (*F_ST_* : fixation index; GO: Gene Ontology; PCA: principal component analysis; RDA: redundancy analysis).

Here, we address this lack of knowledge by proposing a multiplatform, graph-based SV discovery pipeline across numerous genomes (Figure 1A) in order to catalog genetic polymorphism in Romaine and Puyjalon salmon, allowing us to investigate candidate adaptive variation within these populations. With this approach, we primarily targeted small (50 - 1000 bp) to intermediate-sized SVs (< 5kb), <u>as direct SV calling based on short reads and long reads is more accurate and powerful in this range of length (Mahmoud et al., 2019).</u> This study thus served as an unprecedented opportunity to characterize SVs, SNPs and small indels in North American Atlantic salmon, as well as to explore the relative contribution of various forms of genetic variation to fine-scale adaptation and population differentiation.

## 2 Materials and Methods

### 2.1 Sampling, DNA extraction and sequencing

#### 2.1.1 Short reads

Manipulations involving fish were authorized by the Comité de protection des animaux de l’Université Laval (permit number: 2021-783). Adipose fin clips were sampled from 60 wild-born adult salmon raised as broodstock at Université Laval’s Laboratoire de Recherche en Sciences Aquatiques (LARSA) and stored in ethanol until use. The samples comprised 31 Puyjalon (16 males and 15 females) and 29 Romaine (14 males and 15 females) individuals.

Spin column DNA extractions were performed using Qiagen’s DNeasy blood and tissue kit according to the manufacturer’s protocol, with the exception of the elution step, which was done twice per sample with 50 µl of water. DNA quality was assessed by concentration measurement and migration on 1% agarose gel. DNA samples were then diluted to 10 ng/µl and sent to Génome Québec’s Centre d’expertise et de services (Montréal, Canada) for library preparation and whole genome sequencing on an Illumina NovaSeq6000, using four S4 PE150 lanes for an anticipated depth of 16X per sample.

#### 2.1.2 Long reads

Among the 60 fish sampled for whole genome short-read sequencing, four (one male and one female for each population) were used for Nanopore long-read sequencing. In order to provide intact high molecular weight DNA, whole blood was extracted from live fish using EDTA-prefilled syringes, followed by humane euthanasia by decapitation. Blood samples were flash-frozen in liquid nitrogen, transferred to storage tubes and stored at -80°C until use.

High molecular weight DNA extraction was performed twice for each fish using Circulomics’ CBB protocol for nucleated blood (EXT-NBH-001; Circulomics, 2021), from 6 µl of blood mixed with 194 µl of ice-cold PBS. DNA quality was assessed by measuring concentration with Qubit and migrating DNA on a 0.5% agarose gel. DNA samples were then sent to the Centre for Integrative Genomics (CIGENE) at the Norwegian University of Life Science (NMBU) for sequencing. DNA fragments shorter than 25 kb were removed by size selection with Circulomics’ Short Reads Eliminator kit, and seven libraries were prepared for each sample using the SQK-LSK110 kit (Oxford Nanopore Technologies).

Sequencing was performed on a PromethION24 in short serial runs following protocol NFL_9076_v109_revA. Each sequencing run was terminated after a few hours, when the number of active pores dropped to below 10%, in order to recover pores by nuclease-flushing flow cells, which were then refilled with the same DNA preparation for a next short sequencing run. Two FLO-PRO002 flow cells were used for each sample, which were each filled with six and five loadings, respectively, in order to obtain an approximate coverage of 20X. Basecalling was done with Guppy version 5.0.13 (high-accuracy basecalling model) and raw reads were filtered for a minimum qscore of nine. The average yield for the four samples was 47.1 Gb of DNA, while the mean N50 was 39.5 kb.

### 2.2 Characterization of genetic variation

#### 2.2.1 Raw sequencing data preprocessing

##### 2.2.1.1 Short reads

The Ssal_Brian_v1.0 assembly, derived from a North American wild salmon from Newfoundland (Norwegian University of Life Sciences, 2022; GenBank assembly accession: GCA_923944775.1; project accession: CAKLZZ000000000.1), was used as the reference genome for all downstream analyses. This genome features 28 chromosomes with two known polymorphic rearrangements, i.e., the translocation of chromosome ssa01’s p arm (ssa01p) to ssa23 (ssa01-23) (Lehnert et al., 2019), and the fusion of chromosomes ssa26 and ssa28 (Brenna-Hansen et al., 2012).

Raw Illumina data was processed using the wgs_sample_preparation pipeline (https://github.com/enormandeau/wgs_sample_preparation). Adapters and low-quality ends were first trimmed from raw reads by running fastp 0.20.0 (Chen et al., 2018) with default parameters. Trimmed reads were then mapped to the indexed reference genome (samtools *faidx* command, version 1.8; Danecek et al., 2021) using BWA MEM (Li, 2013), allowing a minimum mapping quality of 10 (-q 10). Duplicate reads were filtered out of the alignment with *MarkDuplicates* (Picard 1.119; Broad Institute, 2019). After indexing the resulting bam files with Picard *BuildBamIndex*, mapping was refined around candidate indels using GATK 3.6-0 *RealignerTargetCreator* and *IndelRealigner* (McKenna et al., 2010) and overlapping read pairs were clipped to preserve read regions with the highest average quality using bamUtil 1.0.14 *clip overlap* (Jun et al., 2015). Finally, we used samtools *addreplacerg* to add unique read group names for each sample’s bam file, which is a requirement for some variant calling tools we used.

##### 2.2.1.2 Long reads

Since each sequencing run produced multiple raw read files, all fastq files obtained for a given sample were first concatenated to yield a single fastq file per sample. Raw reads were filtered for an average minimum quality of 10 and a minimum read length of 1,000 bp using NanoFilt 2.0.8 (DeCoster et al., 2018). We mapped filtered reads to the Ssal_Brian_v1.0 assembly with Winnowmap version 2.03 (Jain et al., 2020; Jain et al., 2022) using default parameters and a *k*-mer size of 15 (-k 15). The complete preprocessing pipeline (ONT_data_processing v1.0.0) can be found at https://github.com/LaurieLecomte/ONT_data_processing.

#### 2.2.2 SNP and short indel (1-50 bp) calling

SNPs and small indels were called exclusively from short-read data, as higher basecalling error rates in long-read data are likely to interfere with SNP detection (Rang, Kloosterman & de Ridder, 2018; Ahsan et al., 2021). Variant calling was performed in all 60 samples at once and for each chromosome separately, using bcftools *mpileup* and *call* (version 1.16) and requiring a minimum mapping quality of five at a given site (-q 5). The 28 single chromosome VCF files were then concatenated with bcftools *concat*.

In order to apply the same filtering criteria as SVs (described below), samples without at least four supporting reads and a minimum genotype quality of five for a given variant, or that had more than five times the anticipated whole genome short-read sequencing coverage (80) or an exceedingly high genotype quality (GQ = 127), were assigned the genotype “missing” (“./.”) using bcftools *+set-GT* (version 1.15). Finally, we kept SNPs and small indels that had a minor allele frequency between 0.05 and 0.95 and that were genotyped in at least 50% of samples (i.e., population-scale filters), using bcftools *filter* (version 1.13). The full SNP and indel calling pipeline is available at https://github.com/LaurieLecomte/SNPs_indels_SR (version v1.0.0).

#### 2.2.3 Structural variant calling

##### 2.2.3.1 Short reads

In order to alleviate some of the challenges inherent to SV detection (e.g., low precision), we proposed an ensemble approach where SVs were first called independently with three separate tools, then merged across tools in order to obtain a union callset, which we filtered for calls supported by at least two callers. We assumed that SVs confidently called by multiple tools are more likely to be true positives than SVs called by a single tool.

The three callers used in combination were chosen based on reported performance in previous studies and benchmarks (Cameron, Di Stefano & Papenfuss, 2019; Kosugi et al., 2019; Mérot et al., 2023; Stenløkk, 2023). Each caller was provided with the same 60 bam files, as well as the reference genome used for mapping reads. We first ran DELLY (version 1.1.6; Rausch et al., 2012) following guidelines for germline calling in high coverage genomes (https://github.com/dellytools/delly#germline-sv-calling). Putative SVs were first called separately in all samples and in each of the 28 chromosomes, then merged together in order to obtain a list of known SV sites to be genotyped by DELLY in each sample. Genotyped SVs were then merged across all samples into a unified, multisample VCF. We filtered for deletions (DEL), insertions (INS), duplications (DUP) and inversions (INV) labelled as PASS and PRECISE using bcftools 1.13 *filter*. Next, we used Manta version 1.6.0 (Chen et al., 2015) according to instructions for germline joint samples analysis (https://github.com/Illumina/manta/blob/master/docs/userGuide/README.md#germline-configuration-examples). We parallelized SV calling across the 28 chromosomes instead of across samples, since Manta has no built-in procedure for merging calls across samples. SVs tagged as BND (breakends) were converted into explicit inversions using the script convertInversion.py provided in Manta’s installation directory. The 28 chromosome-specific VCFs were then concatenated into a single multisample file, which was filtered for PASS and PRECISE calls as well. The last short-read-based caller included in the pipeline was LUMPY (Layer et al., 2014), through smoove (version 0.2.7; Pedersen, Layer & Quinlan, 2020). Following recommendations for population-level calling (https://github.com/brentp/smoove#population-calling), we called SVs in the same manner as with DELLY. Only DEL, DUP, INV calls labelled as PRECISE were retained.

The three SV sets were then merged together using Jasmine version 1.1.5 (Kirsche et al., 2023), which integrates various information including chromosome, position, end, size and type to determine whether SV calls from different files or samples refer to the same SV or not. We ran Jasmine with parameters “--mutual_distance --max_dist_linear=0.25”, so that the maximum allowed distance required between two SVs for them to be merged is correlated with their size. The merged VCF was then edited with a custom R script (R version 4.1.2; R Core Team, 2021) in order to convert symbolic alternate alleles to explicit sequences and to standardize VCF fields. The formatted merged VCF was finally filtered for calls supported by at least two callers, with bcftools *filter*. Moreover, in accordance with the most prevalent definition of SVs (Feuk, Carson & Scherer, 2006), variants smaller than 50 bp were considered as small indels instead of SVs and were therefore filtered out. This first set of merged SVs will be referred to as the short-read SV set (SR SVs). The detailed short-read SV calling pipeline can be found at https://github.com/LaurieLecomte/SVs_short_reads (version v1.0.0).

##### 2.2.3.2 Long reads

The SV calling procedure for Oxford Nanopore data is equivalent to the pipeline described above for SV detection from short reads, i.e., independent SV detection with three different tools, merging of SV calls across callers, and filtering for calls supported by at least two tools. However, since most long-read-based SV callers do not support multisample calling, SVs were first called separately for each sample using all three chosen tools, then merged across samples (across-sample merge) to obtain a single VCF per caller. The three multisample VCFs were then merged together (across-caller merge) to obtain the long-read SV set (LR SVs). The pipeline is available at https://github.com/LaurieLecomte/SVs_long_reads (version v1.0.0).

We ran Sniffles 2.0.7 (Sedlazeck et al., 2018a; Smolka et al., 2022) (default settings and “--output-rnames --combine-consensus” options) on each sample and filtered for PASS and PRECISE calls. We then refined alternate allele sequences and breakpoints for insertions, deletions and some duplications by running Iris (Kirsche et al., 2023): we first preprocessed each sample’s VCF with Jasmine (“--dup_to_ins -- preprocess_only”), and ran Iris with parameters “--keep_long_variants --also_deletions”. The four samples’ refined VCFs were merged together using Jasmine --ignore_strand --mutual_distance -- allow_intrasample --output_genotypes, and refined SVs were then converted back to their original type with Jasmine --dup_to_ins --postprocess_only. The multisample Sniffles VCF was finally filtered again for PASS and PRECISE insertions, deletions, duplications and inversions. SVs were also called with SVIM 2.0.0 (Heller & Vingron, 2019) using parameters “--insertion_sequences --read_names -- max_consensus_length=50000 --interspersed_duplications_as_insertions”, following the same procedure as Sniffles to produce a multisample VCF with PASS calls. Last, we used NanoVar 1.4.1 (Tham et al., 2020) with default settings. Supporting reads’ names were added manually using a custom R script in order to allow for the refinement of SV breakpoints by Iris. The three multisample VCFs were finally merged together with Jasmine, formatted with custom R scripts and filtered, as described for the SR SV set.

##### 2.2.3.3 Combination of SV datasets

A final merging step was performed for combining the short-read and long-read SV sets using Jasmine with parameters “--ignore_strand --ignore_merged_inputs --normalize_type --output_genotypes”, resulting in a large union set of candidate SVs to be genotyped in the 60 Atlantic salmon genomes (https://github.com/LaurieLecomte/merge_SVs_SRLR; version v1.0.0).

#### 2.2.4 SV genotyping

We implemented a graph-based genotyping pipeline (https://github.com/LaurieLecomte/genotype_SVs_SRLR, version v1.0.0) using the vg toolkit version 1.46.0 (Hickey et al., 2020), following recommendations from https://github.com/vgteam/vg/wiki/SV-genotyping-with-vg#sv-genotyping-with-vg-call. We first built an indexed reference graph structure from the reference genome fasta and the SV VCF file using *vg autoindex*, then computed snarls, i.e., sites of known variation in the genome graph, with the *vg snarls* command. We then remapped short reads to the variant-aware reference graph for all samples separately using *vg giraffe* (Sirén et al., 2021), computed read support for variation sites (*vg pack*), then genotyped these sites (*vg call*). We used bcftools *+set-GT* on each sample’s VCF to set the genotype as missing (“./.”) for calls that were not supported by at least four reads and that had a quality score lower than five, or that had an extreme quality score (GQ = 256) or an extreme depth (DP = 80), as such calls tend to be false positives (Cameron, Di Stefano & Papenfuss, 2019). All 60 sample VCFs were merged together with bcftools *merge*. The genotyped SV set was finally filtered for variants with a minor allele frequency between 0.05 and 0.95, and less than 50% missing data. <u>The 50% missingness threshold was arbitrary and based on Mérot et al. (2023). Comparison with both less and more stringent missing data proportion thresholds showed that the choice of threshold did not impact the post-filtering variant count differently for SVs than for SNPs or indels (</u>Suppl. Table 1).

In an effort to better link the genomic context and the confidence in SV calling, we compared the frequency distributions of both high-quality SVs that passed all filtering steps and low-quality, filtered out SVs in two genomic features known to interfere with SV calling: highly similar regions resulting from whole-genome duplication events (e.g., syntenic regions) and repeated content (repeats and transposable elements). To identify syntenic regions, we followed the steps described by Dallaire et al. (2023): in summary, we aligned the genome to itself with nucmer (built-in mapper in MUMmer version 4.0.0; Marçais et al., 2018), then performed synteny analysis with SyMAP (Soderlund, Bomhoff & Nelson, 2011), et re-mapped syntenic blocks to the genome with LASTZ (version 1.04.15, Harris, 2007) to get the homology percentage. We identified repeats and transposable elements using RepeatMasker (version 4.0.8). To distinguish between probable false positive SVs and probable true positive SVs, we relied on the filtering criteria we applied on the sample level (read depth and genotype quality) and on the population level (on minor allele frequency and missing data proportion) during the genotyping procedure. Using bedtools *window*, we then extracted excluded and filtered SVs overlapping with either a syntenic region or a repeat region (within a 100-bp window), or both.

In addition, because information on putative SVs is lost at the genotyping step, we applied the procedure described in Supplementary Methods 1 to match genotyped SVs with a known putative SV based on the correspondence of position and allele length, in order to retrieve <u>information on</u> variant type, length and platform support (e.g., short- and/or long-read). <u>This information allowed us to perform additional analyses on the set of genotyped and matched SVs. Indeed, in order to see if long-read SVs could be reliably genotyped from short-read data and a pangenome, we examined the concordance between the genotypes called by vg for long-read SVs and the genotypes called by long-read-based SV callers prior to merging datasets across platforms. First, for the four samples sequenced with both short-and long-read platforms, we extracted the three genotypes (from Sniffles, SVIM and NanoVar) for each long-read SV. We then determined the consensus genotype when possible, e.g., if at least two callers called the same genotype in a given sample. Each SV was labelled as concordant when its consensus genotype matched the corresponding genotype outputted by vg, or as non-concordant if its consensus genotype differed from the vg genotype. Alternatively, when all three callers outputted different genotypes for a given SV in a given sample, no consensus genotype could be inferred, and therefore the concordance between callers and vg could not be determined. We also performed this procedure for short-read SVs for comparison purposes. This concordance analysis is detailed in the scripts compare_GTs_LR_vs_vg.sh and compare_GTs_LR_vs_vg.R from the genotype_SVs_SRLR pipeline.</u>

### 2.3 Population genomics analyses

#### 2.3.1 Differentiation between the Romaine and Puyjalon populations

We used ANGSD version 0.937 (Korneliussen, Albrechtsen & Nielsen, 2014) for performing various population and comparative genomics analyses on SVs, SNPs and small indels separately by adapting a previous pipeline designed for SNPs (https://github.com/clairemerot/angsd_pipeline). To investigate population structure, we first performed principal component analysis (PCA) on a normalized covariance matrix produced from input VCF files using VCFtools (version 0.1.16; Danecek et al., 2011), pcangsd (Meisner & Albrechtsen, 2018) and custom R scripts. From input VCF files, we then estimated average genome-wide fixation index (*F_ST_*; Weir & Cockerham, 1984) from each population’s allele frequency spectrum using ANGSD’s -doSaf and realSFS functions (version 0.937). We also computed *F_ST_* along the genome by sliding windows of 100 kb (per-window *F_ST_*), as well as for each variant (per-variant *F_ST_*).

We employed two complementary approaches for identifying candidate variants likely involved in local adaptation (Figure 1C). We first extracted the most highly differentiated variants falling within the upper 97% per-variant *F_ST_*quantile (*F_ST_* outliers). We also performed Fisher’s exact tests on per-population allelic counts at each site and identified outliers with a corrected *p*-value (*q*-value) lower than 0.01 (Benjamini & Hochberg, 1995). We then extracted common outliers between *F_ST_* and Fisher exact tests to yield a set of strongly differentiated variants used for further analysis.

Second, we ran a redundancy analysis (RDA) on the imputed genotype matrix, with the population as the only explanatory variable using the R package vegan (Oksanen et al., 2022). While *F_ST_*and Fisher’s exact test are more likely to detect outlier loci of large effect, RDA allows identifying covarying markers with individually weak effect that may be involved in polygenic control of phenotypic expression (Rellstab et al., 2015; Forester et al., 2018), as previously documented for life history traits such as age at sexual maturity or growth rate (Sinclair-Water et al., 2020; Debes et al., 2021). We defined RDA candidates as variants with loadings falling over the three standard deviations threshold (Forester, Laporte & Manel, 2018). We thus obtained a set of outlier variants and a set of candidate variants for each of the three variant types studied.

#### 2.3.2 Functional analysis of candidate genomic variation

In order to assess the potential functional impact of candidate variants on life-history trait variation observed in Romaine salmon, we investigated the overlap between variants of interest and known genes. We first annotated the Ssal_Brian_v1.0 assembly using the pipeline GAWN v0.3.5 (https://github.com/enormandeau/gawn) based on the transcriptome of the Ssal_v3.1 assembly (GenBank assembly accession: GCF_905237065.1) and filtered out possible duplicate annotations, which produced a list of 36,697 known genes.

For each set of variants of interest, we ran bcftools *window* (version 2.30.0; Quinlan et al., 2010) to identify a set of overlapping genes located within 10 kb of at least one variant. We then performed Gene Ontology (GO) enrichment analysis on each gene set with goatools 1.2.3 (Klopfenstein et al., 2018), using the list of 36,697 of genes from GAWN as the background (population) set and the go-basic database version 1.2 (2022-07-01; http://release.geneontology.org/2022-07-01/ontology/go-basic.obo). Only enriched terms that referred to a biological process (BP) and with a corrected *p*-value (Benjamini & Hochberg, 1995) under 0.1 were considered significant and preserved. We then used REVIGO (Supek et al., 2011) to cluster significant GO terms by semantic similarity with a cutoff value of 0.5 (“small list”), for easier interpretation. All scripts used for population genomics analyses can be found at https://github.com/LaurieLecomte/SVs_SNPs_indels_compgen (version v1.0.0).

## 3 Results

### 3.1 Long reads revealed more variants while short reads allowed population-scale SV genotyping

SVs were identified through our multistep calling procedure involving both short- and long-read data, where different callers and datasets showed high variability in the number, types and sizes of SVs detected. Indeed, short reads revealed mostly deletions, whereas long reads allowed the detection of many more SVs, especially deletions, insertions and duplications. Among short-read-based callers, Manta reported the most SVs of various types and sizes (151,103), while smoove called the least (28,164), almost exclusively deletions (Table 1 & Suppl. Fig. 1). In total, 238,492 SV calls were merged across the three callers, of which only 15.5% (37,041) were shared by at least two tools and were longer than 50 bp: this short-read SV set primarily consisted of deletions (34,761) smaller than 100 bp, and very few duplications (318) and insertions (849) (Table 2 & Suppl. Fig. 2).

**Table 1.**
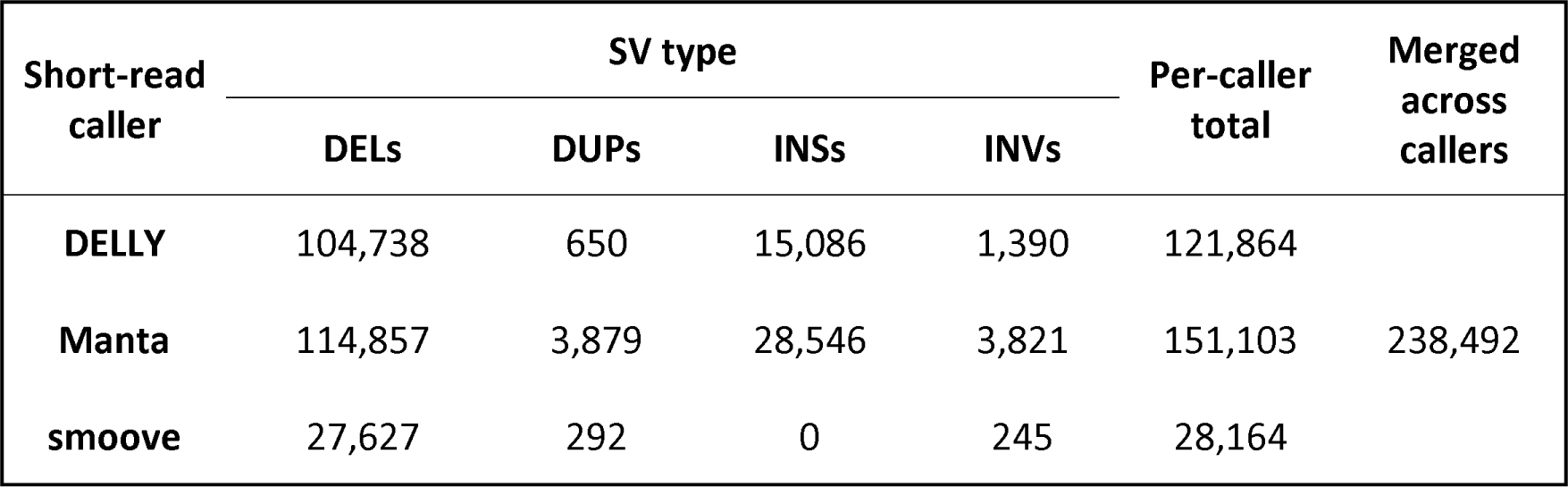
Number of SVs reported by each short-read-based caller, and number of SV calls merged across these callers (DELs: deletions; DUPs: duplications; INSs: insertions; INVs: inversions).

**Table 2.** Number of SVs in the filtered short-read SV set (SR SVs), obtained by merging calls across the three short-reads-based callers, then filtering for a minimum of two supporting tools and a minimum length of 50 bp (DELs: deletions; DUPs: duplications; INSs: insertions; INVs: inversions).

| Short-read caller pair | SV type |  |  |  | Per-pair total |
| --- | --- | --- | --- | --- | --- |
|  | DELs | DUPs | INSs | INVs |  |
| **DELly + Manta** | 7,807 | 108 | 849 | 894 | 9,658 |
| **DELly + smoove** | 2,216 | 122 | 0 | 21 | 2,359 |
| **Manta + smoove** | 1,434 | 26 | 0 | 69 | 1,529 |
| **DELly + Manta + smoove** | 23,304 | 62 | 0 | 129 | 23,495 |
| **Per-type total** | 34,761 | 318 | 849 | 1,113 | 37,041 |

For the long-read pipeline, SVIM called over 3.5 million SVs, mostly insertions and deletions, more than the two other callers combined (Table 3 & Suppl. Fig. 3). The NanoVar callset was the smallest (454,697 SVs) but featured the largest proportions of duplications (31.8%) and inversions (16.4%) (Table 3). The proportion of all merged long-read SV calls (3,832,032) supported by multiple tools was 12.5%, smaller than for the merged short-read SV set. The 345,695 long-read SVs supported by at least two tools and longer than 50 bp retained for subsequent steps were mostly deletions (57.7%) and insertions (40.0%) and included only 208 inversions (Table 4 & Suppl. Fig. 4).

**Table 3.** Number of SVs reported by each long-read-based caller, and number of SV calls merged across these callers (DELs: deletions; DUPs: duplications; INSs: insertions; INVs: inversions).

| Long-read caller | SV type |  |  |  | Per-caller total | Merged across callers |
| --- | --- | --- | --- | --- | --- | --- |
|  | DELs | DUPs | INSs | INVs |  |  |
| **Sniffles** | 260,921 | 635 | 211,588 | 1,233 | 474,377 | 3,832,032 |
| **SVIM** | 1,738,486 | 41,164 | 1,754,905 | 4,914 | 3,539,469 |  |
| **NanoVar** | 178,957 | 144,922 | 123,349 | 7,469 | 454,697 |  |

**Table 4.** Number of SVs in the filtered long-read SV set (LR SVs), obtained by merging calls across the three long-read-based callers, then filtering for a minimum of two supporting tools and a minimum length of 50 bp (DELs: deletions; DUPs: duplications; INSs: insertions; INVs: inversions).

| Long-read caller pair | SV type |  |  |  | Per-pair total |
| --- | --- | --- | --- | --- | --- |
|  | DELs | DUPs | INSs | INVs |  |
| **Sniffles + NanoVar** | 1,831 | 47 | 2,628 | 208 | 4,714 |
| **Sniffles + SVIM** | 107,490 | 38 | 83,797 | 0 | 191,325 |
| **SVIM + NanoVar** | 17,287 | 7,169 | 13,550 | 0 | 38,006 |
| **Sniffles + SVIM + NanoVar** | 72,871 | 451 | 38,328 | 0 | 111,650 |
| **Per-type total** | 199,479 | 7,705 | 138,303 | 208 | 345,695 |

Merging both short- and long-read SVs yielded a set of 361,107 putative SVs, mainly deletions (59.1%) and insertions (38.3%) (Table 5 & Figure 2). The vast majority of these merged SVs, i.e., 89.7%, were exclusively called from long reads, including almost all (99.4%) of the 138,404 insertions identified (Suppl. Table 2). Similarly, 96.0% of duplications and 83.7% of deletions were uniquely detected from long reads. By contrast, only 15,412 SVs, or 4.3% of the merged SV set (Table 5), were unique to the short-read SV callset, including nonetheless 83.3% of all inversions identified (Suppl. Table 2). Moreover, the 21,629 SVs called from both short and long reads accounted for 5.9% of all merged SVs (Table 5), meaning that 58% of all short-read SVs were also supported by long-read data.

**Figure 2.**
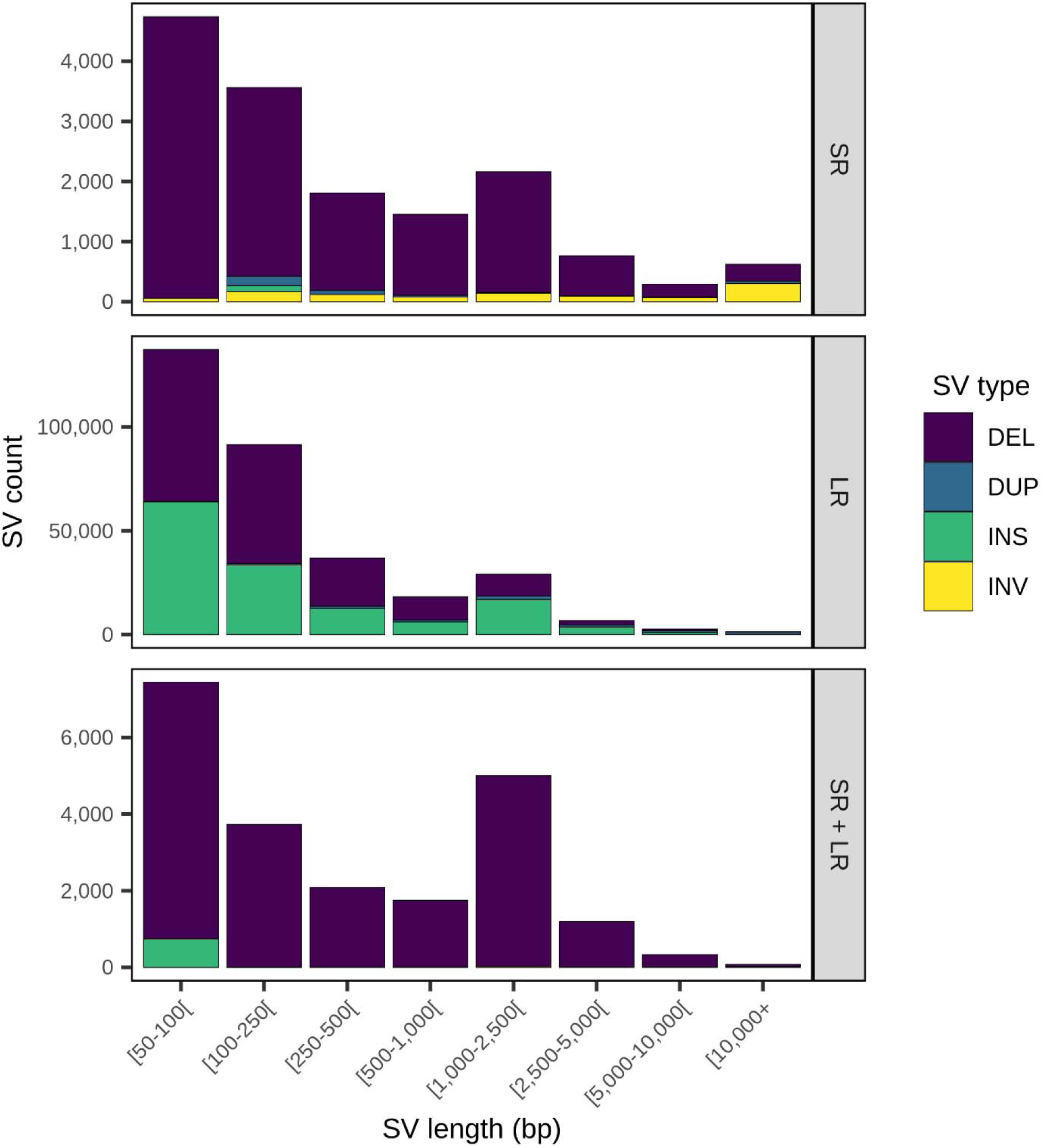
Merged SV count by SV type, length and sequencing platform (SR: short-read; LR: long-read; SR + LR: short- and long-read). These represent putative SVs in the genomes of Romaine and Puyjalon salmon, prior to genotyping (DELs: deletions; DUPs: duplications; INSs: insertions; INVs: inversions).

**Table 5.** Merged SV count by type and sequencing platform (SR: short-read; LR: long-read; SR + LR: short- and long-read). These represent putative SVs in the genomes of Romaine and Puyjalon salmon, prior to genotyping and filtering (DELs: deletions; DUPs: duplications; INSs: insertions; INVs: inversions).

| Platform | SV type |  |  |  | Per-platform total |
| --- | --- | --- | --- | --- | --- |
|  | DELs | DUPs | INSs | INVs |  |
| SR | 13,972 | 305 | 101 | 1,034 | 15,412 |
| LR | 178,690 | 7,692 | 137,555 | 129 | 324,066 |
| SR + LR | 20,789 | 13 | 748 | 79 | 21,629 |
| **Per-type total** | 213,451 | 8,010 | 138,404 | 1,242 | 361,107 |

The merged SVs were represented in a variant-aware genome graph on which we mapped short-read data to genotype SVs in all 60 individuals. On average, 333,031 SVs were genotyped in each sample using the graph-based pipeline, for a total of 344,468 distinct SVs. As expected owing to the complex nature of SVs, the average proportion of missing data per site was over four times larger than for raw SNPs (Suppl. Table 3). 40.8% of raw genotyped SVs did not meet the minimum coverage required in at least half of samples, while two thirds had a minor allele frequency under 5% (Suppl. Table 4). Consequently, filtering on both proportion of missing data and minor allele frequency considerably reduced the SV set to 115,907 high-confidence variants (Table 6), or about 33.6% of the 344,468 raw genotyped SVs. These 115,907 SVs were used for subsequent population genomics analyses.

**Table 6.** Number of filtered SVs, SNPs and small indels used for population genomics analyses along with summary statistics for each variant set. These variants met the population-level filters, i.e., had a minor allele frequency between 0.05 and 0.95 and were genotyped in at least 50% of the 60 samples. The proportion of base pairs covered by variants of a given type was estimated for the whole genome and by 100-kb windows.

| Variant type | Number of variants | Genome base pairs covered | | | $F_{ST}$ | |
| --- | --- | --- | --- | --- | --- | --- |
|  |  | Number | Proportion | Maximum per-window proportion | Genome-wide weighted average | Per-variant maximum |
| SVs | 115,907 | 41,858,002 | 0.0168 | 0.941 | 0.044 | 0.702 |
| SNPs | 8,777,832 | 8,777,832 | 0.0035 | 0.029 | 0.065 | 0.981 |
| Indels | 1,089,321 | 10,316,768 | 0.0041 | 0.102 | 0.079 | 0.949 |

### 3.2 SVs encompassed a more extensive fraction of the genome than SNPs and small indels

While SNPs were the most frequent type of variants, SVs impacted a much higher proportion of genome base pairs, with large heterogeneity along the genome. Indeed, in addition to the final 115,907 SVs, we identified 8,777,832 SNPs and 1,089,321 short indels (Table 6) that met the same filters on minor allele frequency and proportion of missing data. SVs added up to over 41.8 Mb (including insertions), or about 1.68% of total genome length, which was approximately 4.8 times more than SNPs (total length: 8.7 Mb; Table 6) and four times more than small indels (total length: 10.3 Mb; Table 6). Similarly, the proportion of base pairs covered by a given variant type per 100-kb window was much more variable for SVs, reaching as much as 94.0% for some regions (e.g., around a 94.1-kb deletion on chromosome ssa09), whereas the maximum observed proportion of base pairs covered by SNPs was only 2.9% and 10.2% for indels (Table 6 & Figure 3).

**Figure 3.**
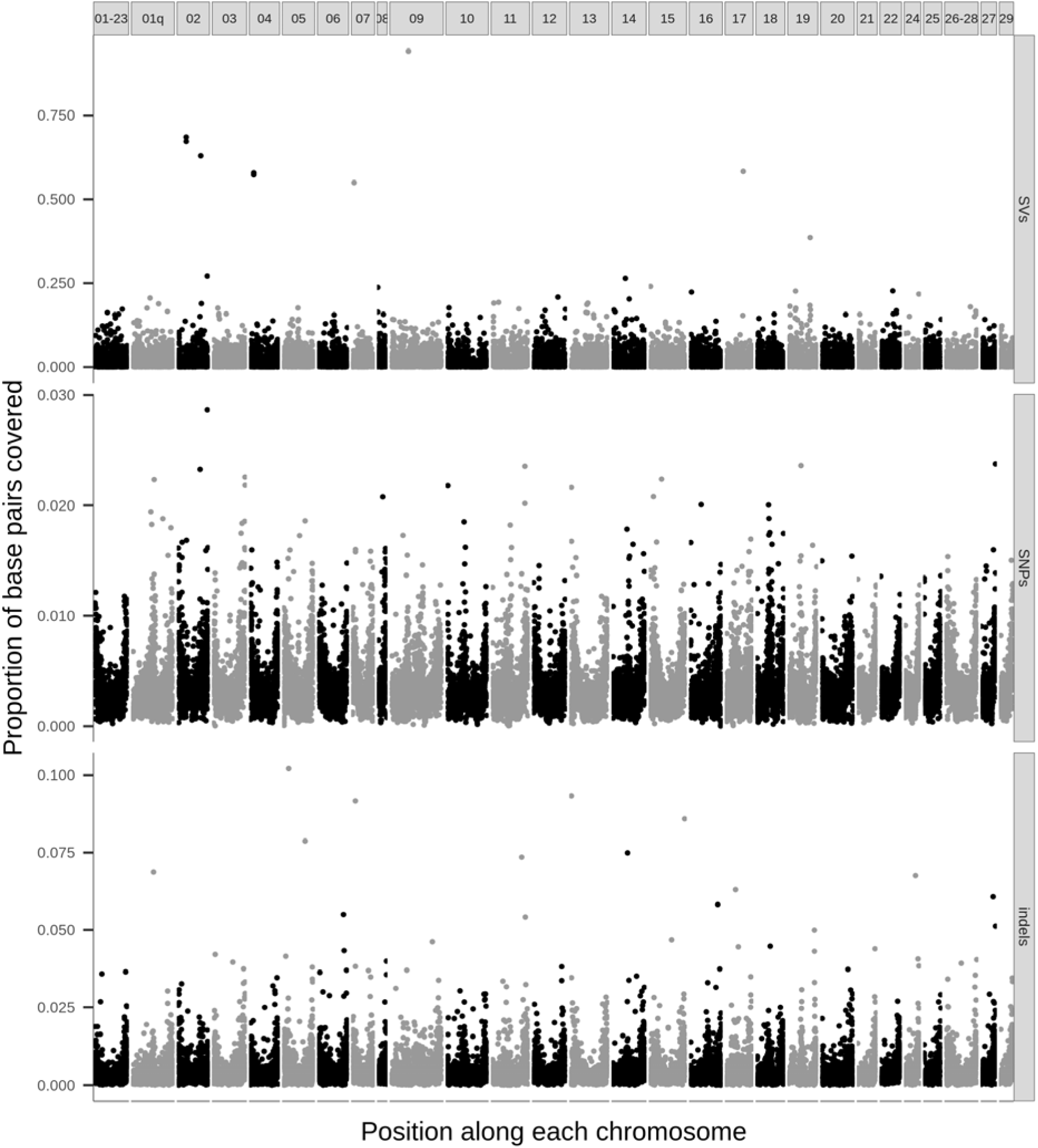
Proportion of base pairs covered by filtered SVs (top), SNPs (middle) and short indels (bottom) per 100-kb windows along the genome. Each vertical panel represents one chromosome.

If we consider the occurrence of variants instead of the number of variable bases, there was no considerable difference in variant density along the genome, as SVs, SNPs and small indels all tended to be more frequent towards the extremities of chromosomes (Suppl. Fig. 5). The gap observed <u>in the number of SVs by 100-kb window (SV density)</u> on chromosome ssa10 could be attributable to a large 2.5-Mb deletion called from short reads, but not successfully genotyped by graphs (Suppl. Fig. 5); this gap was likely not as obvious with SNPs and indels, since numerous markers (over 61,000 SNPs and 8,000 indels) could still be genotyped in samples that did not have this putative deletion.

### 3.3 All genetic variants underlay a congruent population structure

Despite differences in amount and proportion of genome base pairs covered, SVs, SNPs and small indels displayed a consistent population structure and genetic differentiation between the Romaine and Puyjalon populations. Principal component analysis revealed an important differentiation between the two populations, with individual salmon clustering by river along PC1 while PC2 explained variation within Puyjalon samples (Figure 4). This pattern was strongly conserved across all variant types and confirmed anticipated population structure from a previous study (Albert & Bernatchez, 2006). Average genome-wide weighted *F_ST_*values ranged from 0.044 for SVs to 0.065 for SNPs and 0.079 for small indels (Table 6). This observation, along with the strong clustering of samples in principal component analysis, indicates moderate levels of differentiation between Romaine and Puyjalon samples.

**Figure 4.**
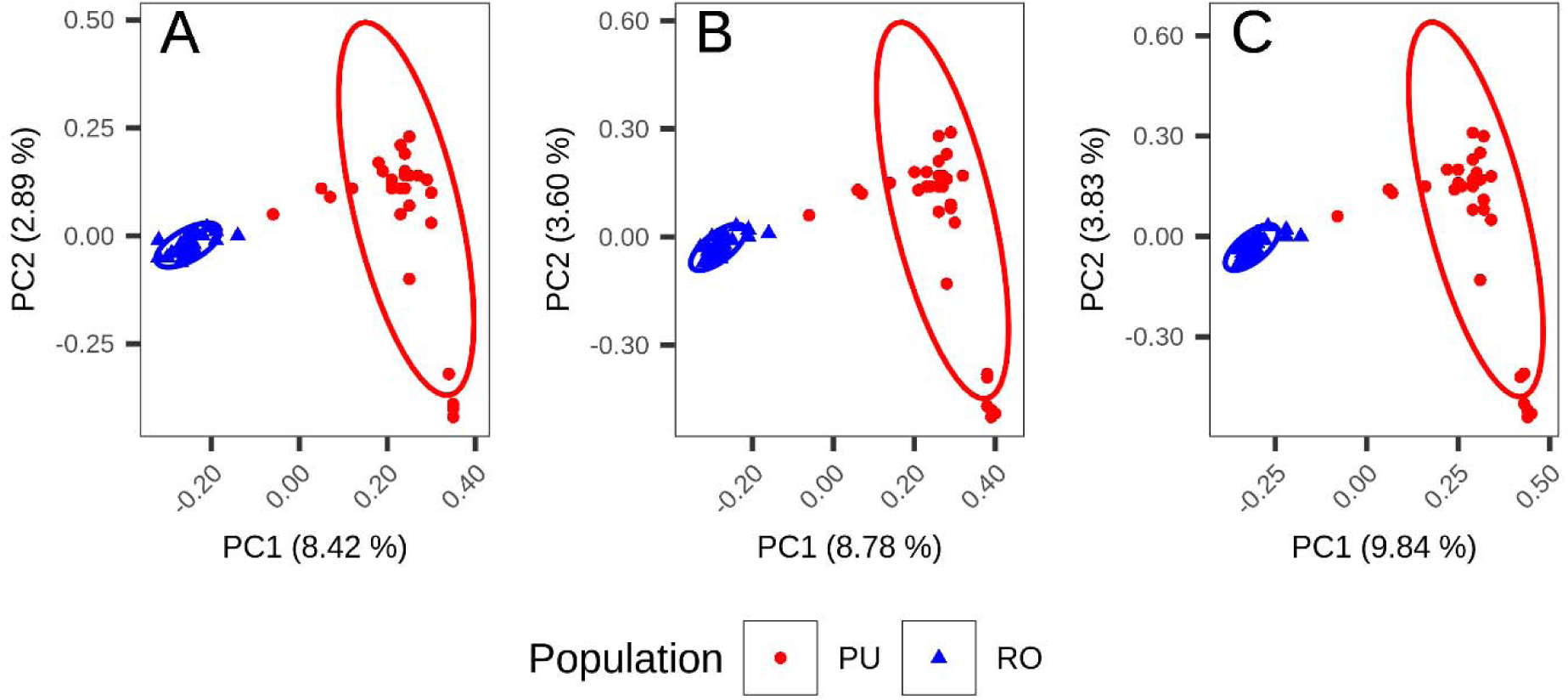
Population structure of Romaine (RO) and Puyjalon (PU) salmon based on principal component analysis from filtered and genotyped **A)** SVs, **B)** SNPs and **C)** short indels. Each point represents one of the 60 salmon sampled for the study.

Differentiation along the genome was also highly variable, as regions of strong per-window *F_ST_* (> 0.2) were numerous and dispersed on all chromosomes (Figure 5). These high *F_ST_* peaks were overall consistent across SVs, SNPs and short indels. However, the correlation was the greatest between SNPs and indels (*R^2^* = 0.930, *p*-value < 0.001), which shared multiple peaks and tended to have higher *F_ST_* than SVs. SVs displayed a weaker correlation with SNPs (*R^2^* = 0.612, *p*-value < 0.001) and indels (*R^2^*= 0.595, *p*-value < 0.001). Only a few per-window *F_ST_*peaks were unique to SVs (e.g., on chromosomes ssa01q, 10, 29; Figure 5). Per-variant *F_ST_* distribution also differed between variant types: values ranged from 0 to 0.981 for SNPs, whereas the maximum per-variant *F_ST_*observed for SVs was 0.702 (Table 6).

**Figure 5.**
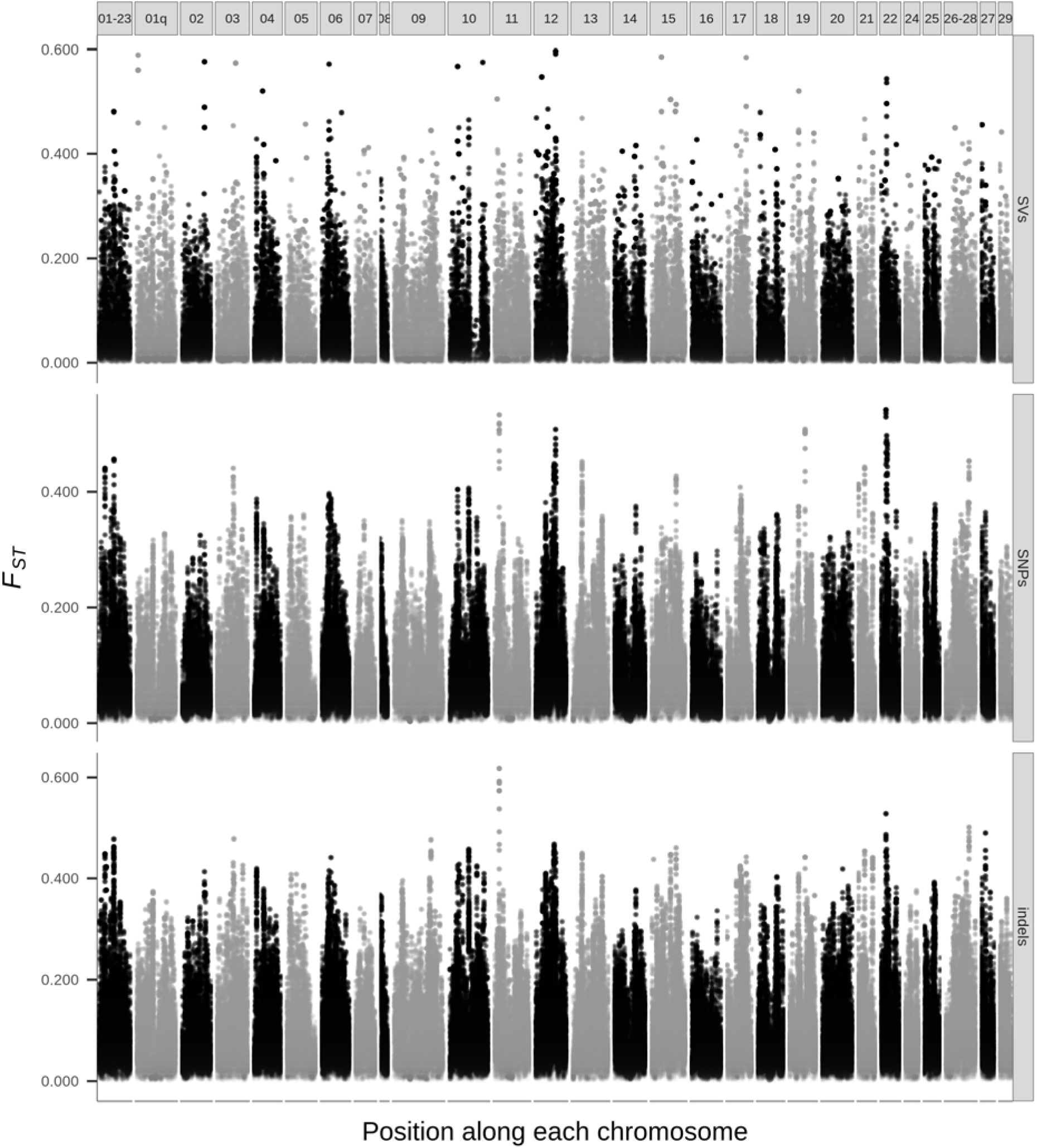
*F_ST_*along the genome between Romaine and Puyjalon salmon, estimated by 100-kb windows (with 10-kb steps) from SVs (top), SNPs (middle) and short indels (bottom). Dotted white lines correspond to the weighted mean genome wide *F_ST_* for each variant type. Each vertical panel represents one chromosome.

### 3.4 Candidate variants for local adaptation overlapped genes involved in putatively important biological functions

Among all filtered variants of each type, we identified those that showed a possible relevance in the putative local adaptation of Romaine and Puyjalon salmon. For each variant type, we reported the most strongly differentiated variants between both populations, as these could include major-effect variants, as well as RDA candidates, which might instead reveal multiple, small-effect loci. We identified 1.62 times more outliers of differentiation than RDA candidates for SVs, whereas SNPs and small indels showed the opposite trend, with 1.59 and 2.23 times more RDA candidates than outliers for SNPs and indels, respectively (Table 7). <u>These outlier and RDA candidate variants did not have more missing data than the others, non-outlier and non-candidate variants (</u>Suppl. Fig. 6<u>), meaning that the apparent differentiation and multilocus signals were not artificially driven by missing genotypes.</u> For each of these six sets of variants of interest, we reported a set of overlapping genes within 10 kb, ranging from only 940 genes for RDA candidate SVs to 15,711 genes for RDA candidate SNPs (Table 8). GO enrichment analysis performed on each of these six gene sets revealed various biological processes, with a redundancy of terms associated with cellular structure and nervous system function. The 1,407 genes located near SV outliers were enriched for 108 terms mostly related to cellular adhesion and junction, as well as to synapse organization (Suppl. Table 5), whereas RDA candidate SV genes were enriched for only 27 GO terms clustered under “chemical synaptic transmission” (Suppl. Table 6). Many of the GO terms that were enriched for outlier SV genes, such as cell adhesion and junction, synapse organization or developmental processes, were also among the 223 enriched terms for outlier SNP genes (Suppl. Table 7). A much wider range of biological functions were overrepresented in the 528 GO terms associated with RDA candidate SNP genes, including growth and nervous system development (Suppl. Table 8). Finally, genes that overlapped with outlier and RDA candidate indels were enriched for fewer GO terms (212 and 292, respectively), but enriched terms themselves were similar to those reported for the outlier and RDA candidate SNP gene sets (Suppl. Table 9; Suppl. Table 10). Raw GO enrichment result tables (prior to simplification with REVIGO) can be found in Supplementary Data 1.

**Table 7.** Outlier and candidate variant sets for each type of polymorphism. The intersection outlier set corresponds to variants that were among the upper 3% quantile of per site *F_ST_*and that had a Fisher test *q*-value lower than the 0.01 threshold, whereas RDA candidate variants were identified through RDA (redundancy analysis) using a threshold of three standard deviations.

| Variant type | Outliers |  |  | RDA candidates |
| --- | --- | --- | --- | --- |
| | $F_{ST}$ | Fisher | Intersection | |
| SVs | 3,484 | 5,298 | 2,079 | 1,280 |
| SNPs | 263,316 | 591,621 | 71,953 | 114,637 |
| Indels | 32,544 | 103,829 | 6,223 | 13,871 |

**Table 8.** Number of genes overlapping intersection outlier variants and RDA candidate variants for each type of polymorphism, along with the number of significant enriched GO terms for each gene set. Variants and genes were considered as overlapping if they were located within 10 kb of each other, either from their start or their end positions.

| Variant type | Outliers of differentiation |  |  | RDA candidates |  |  |
| --- | --- | --- | --- | --- | --- | --- |
|  | Variants | Nearby genes | Significant enriched GO terms | Variants | Nearby genes | Significant enriched GO terms |
| SVs | 2,079 | 1,407 | 108 | 1,280 | 940 | 27 |
| SNPs | 71,953 | 11,578 | 223 | 114,637 | 15,711 | 528 |
| Indels | 6,223 | 3,285 | 212 | 13,871 | 5,872 | 292 |

## 4 Discussion

Structural variants now appear as major components of genetic polymorphism with increasingly recognized implications for phenotype and adaptation, but they are inherently difficult to characterize, especially at the population level. By proposing a hybrid pipeline combining short- and long-read sequencing with graph-based genotyping, we aimed to alleviate some of the challenges hampering the study of SVs in population genomics. Applying this pipeline to a set of 60 Atlantic salmon genomes allowed us to catalog a wide spectrum of polymorphism, from SNPs to multi-kilobase SVs. From this catalog, we described the population structure of salmon populations inhabiting two adjacent tributaries, the Romaine and Puyjalon rivers, estimated the level of differentiation between them, and uncovered putative variants underlying local adaptation.

### 4.1 Multiplatform variant detection and pangenome-based approaches facilitate population-scale analysis of SVs

The combination of long- and short-read sequencing with graph-based genotyping is a promising, yet incomplete solution to the challenges raised by analyzing SVs at the population scale. Our results indicate that long reads are a highly valuable asset for SV characterization, because the vast majority of SVs were identified from the only four genomes sequenced in Oxford Nanopore long reads, despite high-coverage (16X) paired-end Illumina short-read data being available for all 60 samples. Long-read data was particularly crucial for detecting putative insertions and characterizing their sequence, as expected from previous studies (Ho, Urban & Mills, 2020). Since insertions and deletions are the most prevalent forms of SVs in genomes (Gamazon & Stranger, 2015), the incorporation of third generation sequencing enabled a considerable gain in the amount of genetic variation uncovered. However, elevated costs remain an obstacle for population-scale implementation of long-read sequencing, especially for species with large genomes such as salmonids, with prices ranging from 22 to 200 $ USD per gigabase depending on technology used (De Coster, Weissensteiner & Sedlazeck, 2021). Here, graph-based genotyping allowed us to exploit short reads to genotype putative SVs in a much higher number of samples whose genomes were only sequenced in short reads. Indeed, about 93.9% of the 115,907 final, filtered and genotyped SVs were initially called from long reads (or from both short and long reads; Suppl. Table 11) and could be genotyped in more than half of all samples, indicating that even though short reads are suboptimal for SV detection, they are highly relevant for genotyping SVs in populations. The hybrid approach thus offered a major increase in overall SV genotyping power at a much lower cost, which therefore represents a reasonable compromise between cost and efficiency.

This multiplatform approach nevertheless does not address all issues pertaining to SV characterization. First, merging SV calls across tools relies on arbitrary thresholds and may be suboptimal, depending on the precision of SV characterization procedures. We made the arbitrary decision to retain only SVs that were supported by at least two callers, which might be an overly conservative filtering approach. Given the high false discovery rates previously reported for SV calling in the Atlantic salmon genome (Bertolotti et al., 2020), we chose to attempt improving precision over sensitivity. However, multitool calling is not guaranteed to improve precision nor sensitivity, depending on datasets and callers used in combination, and callers relying on the same evidence types are likely to call the same false positives (Chaisson et al., 2019; Kosugi et al., 2019; Mahmoud et al., 2019).

In addition, despite using Jasmine, an up-to-date graph-based minimum spanning forest algorithm that integrates numerous SV parameters, a surprisingly small number of all SVs called by multiple tools could be merged together. This is especially true for merged long-read SVs, for which the proportion of shared calls was even smaller than in the merged short-read SV set. We suggest that the lower basecalling accuracy inherent to third generation sequencing data might increase breakpoint imprecision (Lemay et al., 2022) and lead to undermerging of shared variants, especially for those that could not be refined by Iris. For example, long-read-based callers each reported a few thousand inversions, but only 208 were successfully merged across Sniffles and NanoVar. This likely results from imprecision of inversion breakpoints (Sudmant et al., 2015a), as the breakpoints refinement tool (Iris) cannot process inversions at this time, or from differences in how inversions are identified or reported by different tools. All these issues highlight the need for more efficient validation methods for large SV datasets. Since visualization remains the most robust SV validation method (Spies et al., 2015), emerging machine learning-based <u>and/or automated</u> methods show promising applications for this purpose, such as samplot-ML (Belyeu et al., 2021), MAVIS (Reisle et al. 2019) and DeepSVFilter (Liu et al., 2020), but also for SV detection and genotyping (Cue; Popic et al., 2023). <u>We can expect that such tools, which currently primarily target short-read data, will eventually allow automated long-read SV validation as well, thus further improving SV analysis in the upcoming years</u>.

Although graph-based genotyping has been essential for genotyping long-read SVs from short-read data, it is not immune to bias in SV characterization. We explicitly treated genotypes with insufficient read support (or genotype quality) as missing data, and up to 40% of SVs were filtered out due to missing genotypes in at least half of the samples, meaning that very few reads could be mapped to these SV regions in the pangenome. <u>Since long-read SVs had a consistently higher proportion of missing genotypes than short-read SVs, both before and after filtering on genotype quality, depth and minor allele frequency (</u>Suppl. Fig. 7<u>), we speculate that some short reads still cannot be accurately mapped to certain SV regions where long reads could be confidently aligned. This might also explain the lower concordance between the genotypes outputted by vg and the genotypes provided by caller genotypes for candidate long-read SVs than for candidate short-read SVs (</u>Suppl. Table 12<u>). As stated above, the higher breakpoint imprecision for long-read SVs might also increase noise around SV positions and therefore contribute to the poorer mapping of short reads to the graph.</u> Various features of the Atlantic salmon genome are known to promote spurious mapping of reads to the linear reference genome, such as residual tetrasomy (10 to 20%; Houston & Macqueen, 2019), highly similar duplicated regions (81 – 89%; Davidson et al., 2010) and a large proportion of repeats (50 – 60%; de Boer et al., 2007). We can expect that these features also impact pangenome-based mapping and genotyping to some extent, especially repeats (Chen et al., 2019; Outten et al., 2021). <u>Indeed, low-confidence SVs that were filtered out after genotyping were more prevalent in regions with repeated contents (transposable elements and repeats) and/or in syntenic regions with elevated levels of homology following a past whole-genome duplication event in salmonids (</u>Suppl. Fig. 8). Second, very large putative SVs spanning considerable chromosomal regions, such as the 2.5-Mb deletion on chromosome ssa10, could not be successfully genotyped using graphs, likely because such large rearrangements cannot be reliably represented by complex graph structures (Hübner, 2022). This limitation is particularly problematic for the study of SVs in the context of population genomics, as larger rearrangements were found to play a key role in adaptive processes (Wellenreuther & Bernatchez, 2018; Wellenreuther et al., 2019). <u>Alternatively, some of the very large candidate SVs that were not successfully genotyped could have been false positives, as over half of putative SVs larger than 30 kb were inversions and deletions exclusively supported by short-read callers (</u>Suppl. Table 13). Our study would therefore benefit from the addition of complementary approaches, such as assembly comparison or chromatin conformation data, to identify, <u>validate</u> and genotype large SVs (Mérot et al., 2020), hence further expanding the range of SVs identified. Despite these limitations, the multiplatform strategy developed for this study represents a considerable improvement over short-read-only approaches, and the incorporation of novel automated curation approaches will undoubtedly lead to further advancements in population-scale SV characterization.

### 4.2 SVs are a key feature of the Atlantic salmon genome

Our findings showed that the contribution of SVs to standing genetic polymorphism is important in Atlantic salmon. High-confidence, genotyped SVs accounted for 4.8 times more genome base pairs than SNPs. This proportion is in the same order of magnitude as previously estimated using an equivalent approach in lake whitefish, a closely related species (e.g., five times; Mérot et al., 2023). The number of SVs identified in our study is over seven times larger than previously documented in rainbow trout (almost 14,000 SVs; Liu et al., 2021) and in Atlantic salmon (over 15,000 SVs; Bertolotti et al., 2020) in previous studies involving more samples, but relying only on short-read data. Similarly, we reported between 20 and 30 SVs per 100-kb window, whereas the median per-megabase SV count reported by Bertolotti et al. (2020) is under 10. SV counts described in our study might be inflated due to a certain number of false positives in our dataset, since we did not exclude calls located in problematic regions (e.g., high coverage regions, assembly gaps and low complexity regions) nor performed manual curation of SVs. However, such an important difference in SV count can most likely be explained by the integration of long-read sequencing data. Indeed, in the human genome, over six times more high-confidence SVs were identified from long reads (27,662 SVs; Chaisson et al., 2019) than from short reads in another study (4,442 SVs; Abel et al., 2020).

### 4.3 SVs are informative markers relevant for population genomics studies

SVs also appear to reliably capture population structure and differentiation to the same extent as SNPs. The very high correspondence of population structure inferred from PCA across variant types was also observed in previous studies of SVs in soybean (Lemay et al., 2022), in cacao (*Theobroma cacao*; Hämälä et al., 2021), in grapevine (*Vitis vinifera* ssp. *Sativa*; Zhou et al., 2019), lake whitefish (Mérot et al., 2023) and *Corvus* genus species (Weissensteiner et al., 2020). Patterns of fluctuations in per-window variant density and *F_ST_*along the genome were also strongly conserved e.g., regions of high SV density were usually dense in SNPs and short indels as well. <u>We reported a quick linkage disequilibrium decay in all pairs of variants, with minimal linkage between SNPs and SVs for distances greater than 250 bp (</u>Suppl. Fig. 9<u>). This suggests that the observed correspondence between all three types of variants might not be attributable to strong physical linkage between them, but rather that SVs, SNPs and short indels may be subject to similar evolutionary processes in the Romaine and Puyjalon system, despite them</u> <u>being very different in size.</u> Per-variant and per-window *F_ST_* was usually slightly lower for SVs than for SNPs and small indels, which was also reported in the lake whitefish study (Mérot et al., 2023). We suspect that this slight discrepancy is attributable to the fact that the *F_ST_* calculation relies on fewer markers for SVs, thus introducing more noise in *F_ST_* estimates than with SNPs and small indels.

By contrast, in American lobster, a non-related and less structured marine species (Kenchington et al., 2009; Benestan et al., 2015), copy number variants harbored stronger interpopulation differentiation than SNPs and a more defined population structure, correlated with environmental variables (Dorant et al., 2020). Similarly, deletions showed stronger spatial population structure and were under stronger selection than duplications in human populations (Sudmant et al., 2015b). In European starling (*Sturnus vulgaris*), SVs and SNPs revealed different patterns of population structure, interpopulation genetic diversity and divergence across the genome (Stuart et al., 2023). Consequently, we cannot assume that SVs, SNPs and small indels are interchangeable and equally informative in all systems and species as we observed in the Romaine and Puyjalon system. We therefore argue that we ought to characterize SVs in population genomics studies to the same extent as SNPs, as they may display different signatures and provide relevant insights on evolutionary and adaptive processes shaping population structure.

### 4.4 Genetic divergence between the Romaine and Puyjalon populations is likely driven by divergent selection

Given their spatial proximity and habitat overlap, gene flow is expected to occur between the Romaine and Puyjalon populations. The moderate average genome-wide *F_ST_* values estimated from the different classes of variants, as well as a previous estimation based on microsatellites (Albert & Bernatchez, 2006), are consistent with a rate of two to seven migrants per generation, based on Wright’s approximation (1978). However, we reported numerous outliers of differentiation and peaks of very strong *F_ST_* dispersed throughout the genome that are seemingly resisting such gene flow. The persistence of pronounced divergence in these regions could be explained by a few alternative and not mutually exclusive mechanisms. Genetic drift could lead to random differences in variant allelic frequency in both populations. Alternatively, given that recombination rates are known to differ within species and even within populations (Kong et al., 2010; Ritz, Noor & Singh, 2017), some localized regions of low recombination could have emerged independently in both populations, capturing different alleles and being subject to increased genetic drift as a result of apparent reduced effective population size (*N_e_*). Such “differentiation islands” can result from the interplay of variation in recombination rate due to genetic architecture (e.g., the presence of SVs or large rearrangements) and natural selection (Wolf & Ellegren, 2017). Finally, variants with a functional impact could be subject to divergent selection between habitats leading to differences in allelic frequencies.

Our findings tend to support the hypothesis of local adaptation. First, we reported a repeated enrichment for GO terms related to nervous system function for genes nearby outlier and RDA candidate variants, regardless of variant type. We initially expected enrichment for functions related to other observed phenotypic trait variation in the Romaine and Puyjalon system, such as growth, sexual maturation and reproduction. On the contrary, we observed enrichment mainly related to nervous functions. We hypothesize that enrichment for nervous functions could be linked to variation in these traits through their link with age at smoltification that differ between these two populations. Indeed, changes in photoperiod, which are recognized as the main factor triggering smoltification (Hoar, 1988), are perceived and processed through the light-brain-pituitary axis, inducing the hormonal cascade responsible for physiological, morphological and behavioral changes underlying smolt-to-parr transition (Stefansson et al., 2008). Smoltification itself causes reorganization of nervous connections, both at the structural and the chemical level (Ebbesson et al., 2003). Although empirical evidence is required to support this hypothesis, polymorphism around genes involved in nervous system function, development and plasticity could alter the expression or function of these genes and lead to physiological differences underlying variation in age at smoltification and other relevant life history traits in Romaine and Puyjalon salmon. This indirect, but plausible link between observed phenotypic variation and genetic polymorphism does not support the hypothesis of persistent genetic differentiation due to genetic drift alone. Moreover, peaks of differentiation are unlikely a result of low recombination alone because such peaks and outliers of differentiation are dispersed throughout chromosomes and across the whole genome, and not clustered into contiguous and localized regions. Along with preliminary knowledge of the Romaine and Puyjalon system, our results suggest that the persistence of localized regions of strong differentiation could at least partly be attributable to local adaptation in response to divergent selection. Indeed, both rivers differ in habitat quality, substrate and hydrological profiles (Schieffer,1975; Fontaine et al., 2000; GENIVAR, 2002; Belles-Isles et al., 2004; WSP Global, 2019), which may impose different constraints on salmon and thus favor alternate life strategies in both populations.

Besides showing variation in age at smoltification, Romaine and Puyjalon salmon also differ in regards to age at sexual maturity in a controlled hatchery environment (Therrien et al., 2017; Langlois-Parisé et al., 2018; T. Dion et al., 2020; T. Dion, Langlois-Parisé & Proulx, 2020). Interestingly, we found no variant of interest overlapping with major-effect loci previously associated with life history variation in age at maturity in wild and domesticated European salmon populations, such as *vgll3* (Ayllon et al., 2015; Barson et al., 2015; Czorlich et al., 2018) and *six6* (Sinclair-Waters et al., 2020; Waters et al., 2021). While one study in North America highlighted a correlation between *vgll3* polymorphism, sea age and sex in the Trinité river population (Kusche et al., 2017), located in the same geographic region as the Romaine and Puyjalon rivers, other studies did not reveal a significant association between polymorphism in these major-effect loci and age at maturity in other North American populations (Boulding et al., 2019; Mohamed et al., 2019). The genetic architecture of such complex life history traits is possibly variable across populations, especially between highly divergent populations from different continents. In addition, age at maturity was found to have a mixed genetic architecture in both North American-derived farmed salmon (Eisbrenner et al., 2014; Mohamed et al., 2019) and European-origin salmon (Sinclair-Water et al., 2020), involving both major-effect loci and multiple small-effect loci. Since we identified numerous candidate small-effect variants through RDA, we propose that phenotype variation in age at maturity in Romaine and Puyjalon salmon might have a polygenic basis as well.

Further work is required to understand the genetic architecture of major life history trait variation in Atlantic salmon populations as well as other salmonid species. Such work would considerably benefit from an improved knowledge of the full spectrum of genetic variation segregating in populations, especially SVs. The pipelines developed and optimized for this study may therefore contribute this knowledge by facilitating population-scale characterization of SV, <u>as well as serve as a basis for further refinement of variant calling and genotyping procedures in the near future. With ongoing and rapid developments in computational genomic approaches, such as pangenome-based tools or machine-learning-based variant detection and validation, SV analysis is bound to take a significant leap towards robust and reliable characterization in the upcoming years, which will foster their inclusion in evolutionary genomics.</u>

## Supporting information

Raw GO enrichment analysis results

