## Supplementary figures and tables 1 to 4 and 11 to 13 for "Investigating structural variant, indel and single nucleotide polymorphism differentiation between locally adapted Atlantic salmon populations"

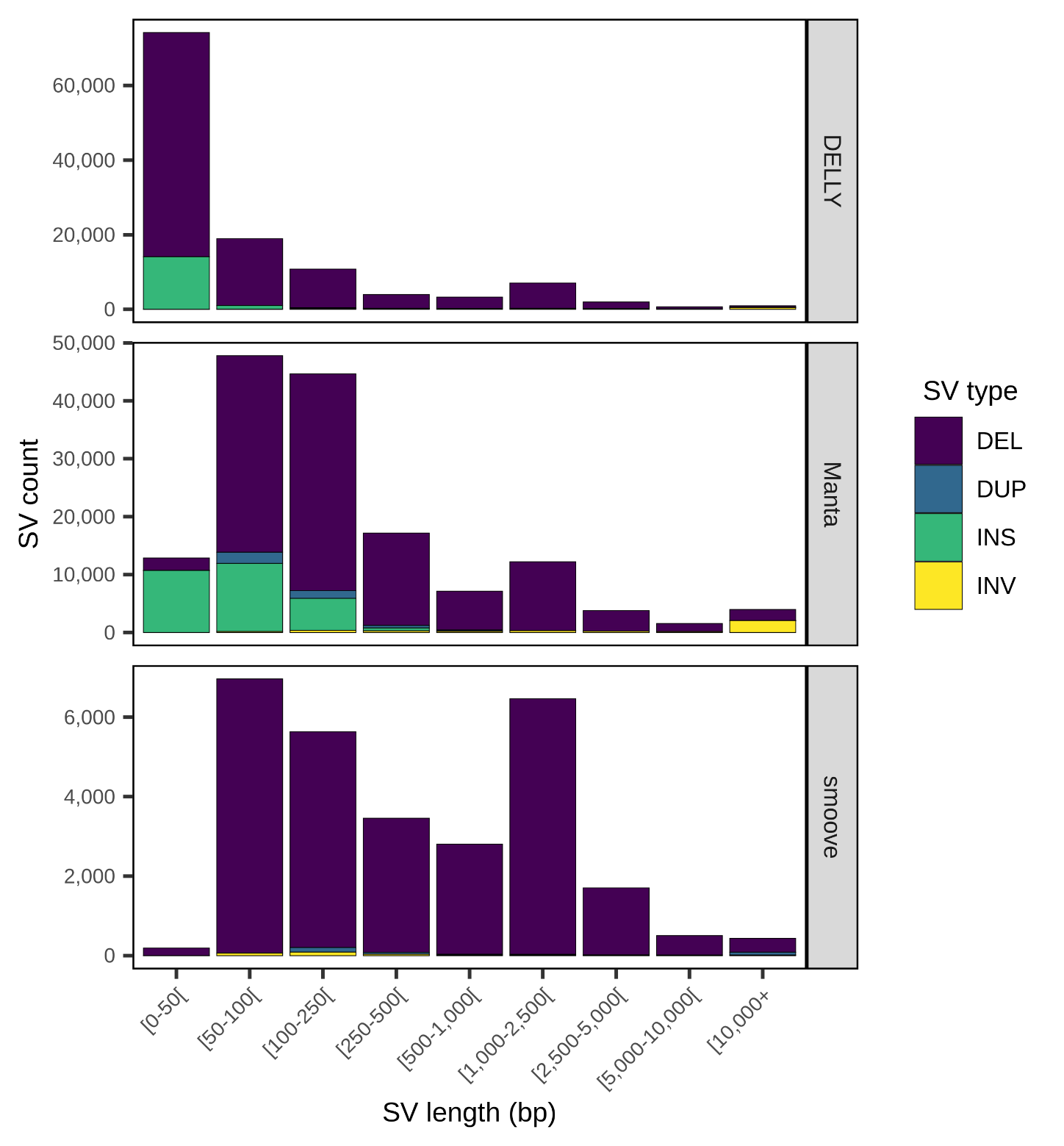

**Supplementary Figure 1.** Number of raw SV calls reported by each short-read-based caller, by type and length (DEL: deletions; DUP: duplications; INS: insertions; INV: inversions).

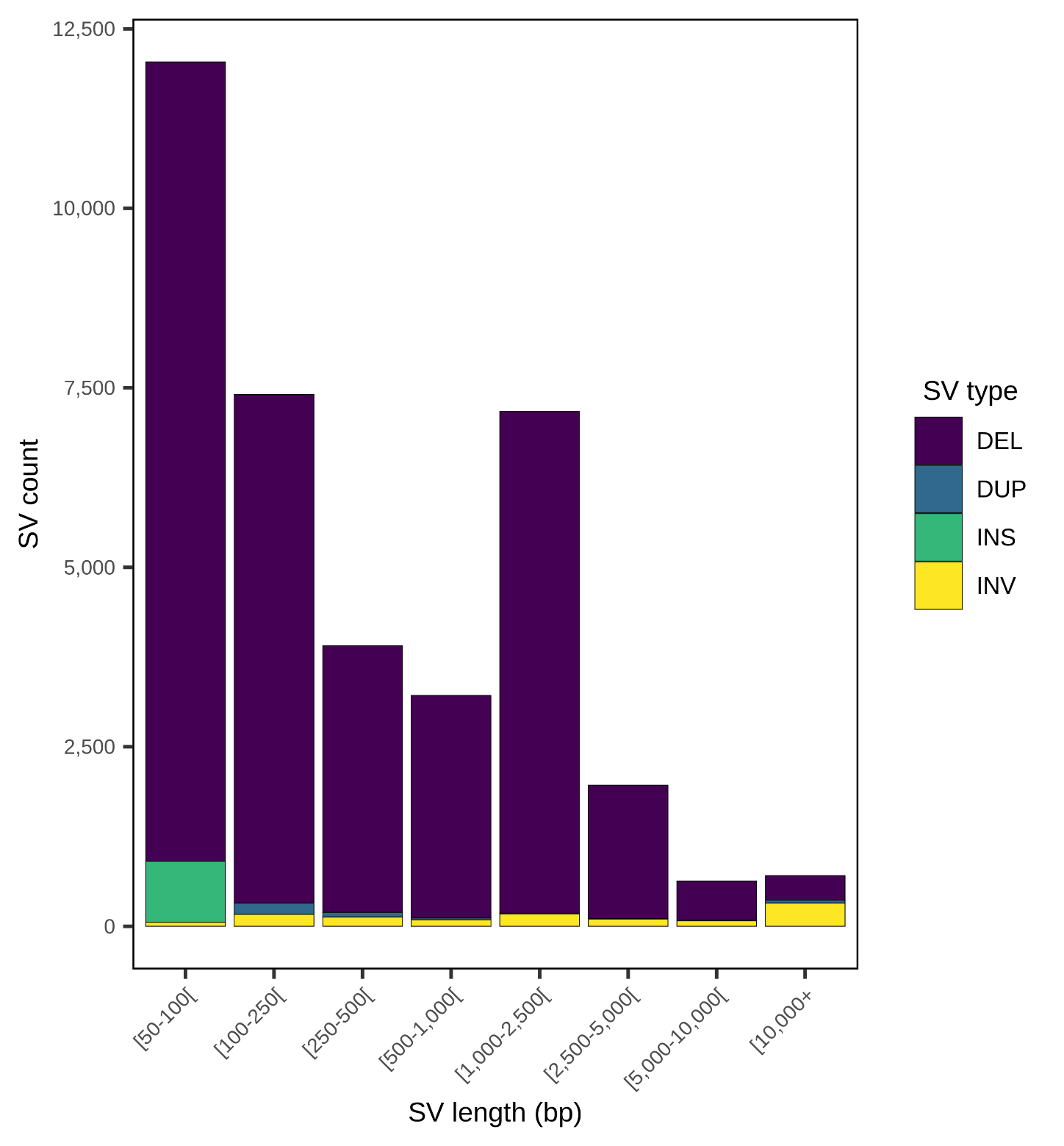

**Supplementary Figure 2.** Number of SVs in the filtered short-read SV set (SR SVs) by type and length, obtained by merging calls across the three short-read-based callers (DELLY, Manta and smoove) and filtering for a minimum of two supporting tools and a minimum length of 50 bp (DEL: deletions; DUP: duplications; INS: insertions; INV: inversions).

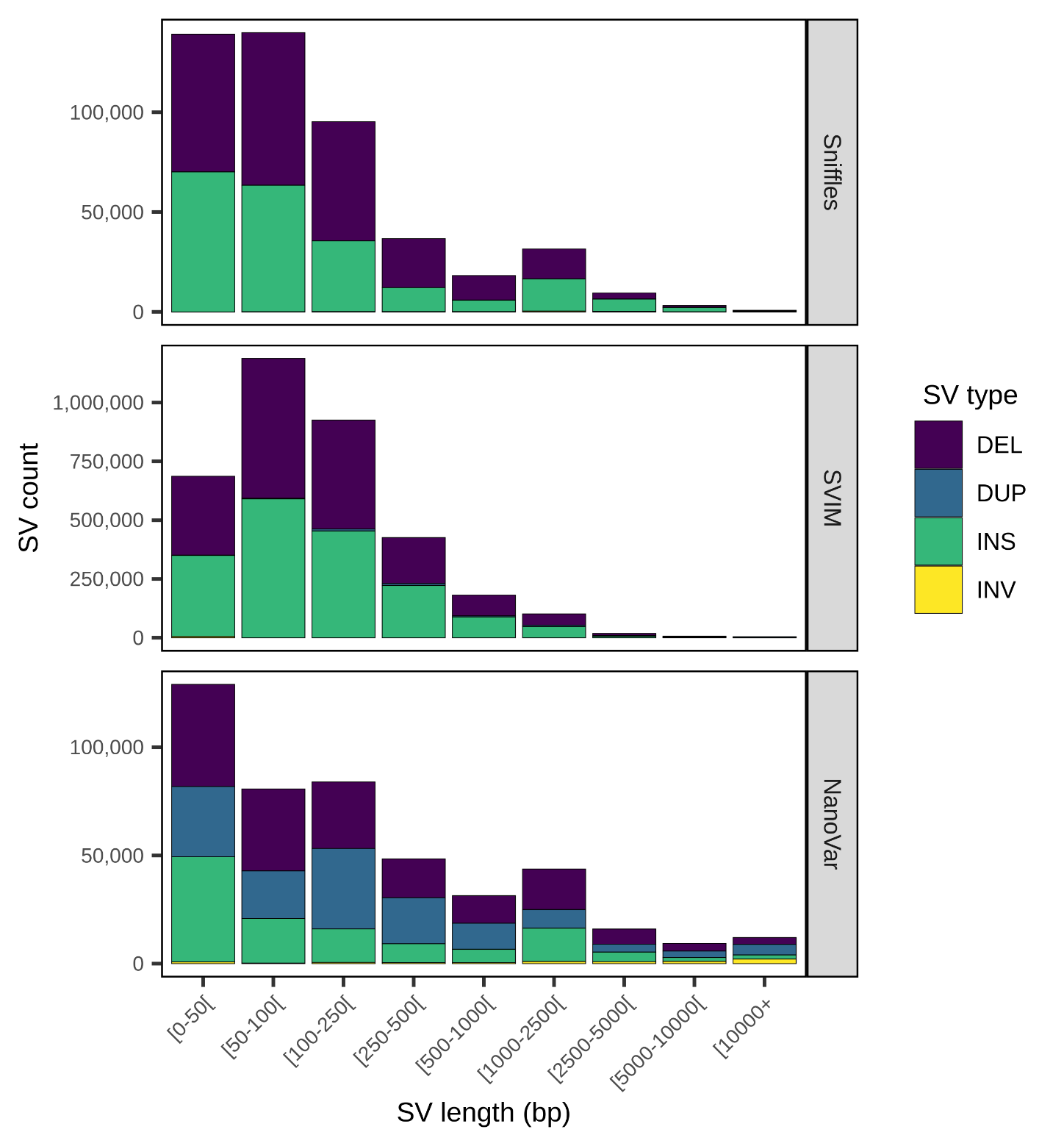

**Supplementary Figure 3.** Number of raw SV calls reported by each long-read-based caller by type and length (DEL: deletions; DUP: duplications; INS: insertions; INV: inversions).

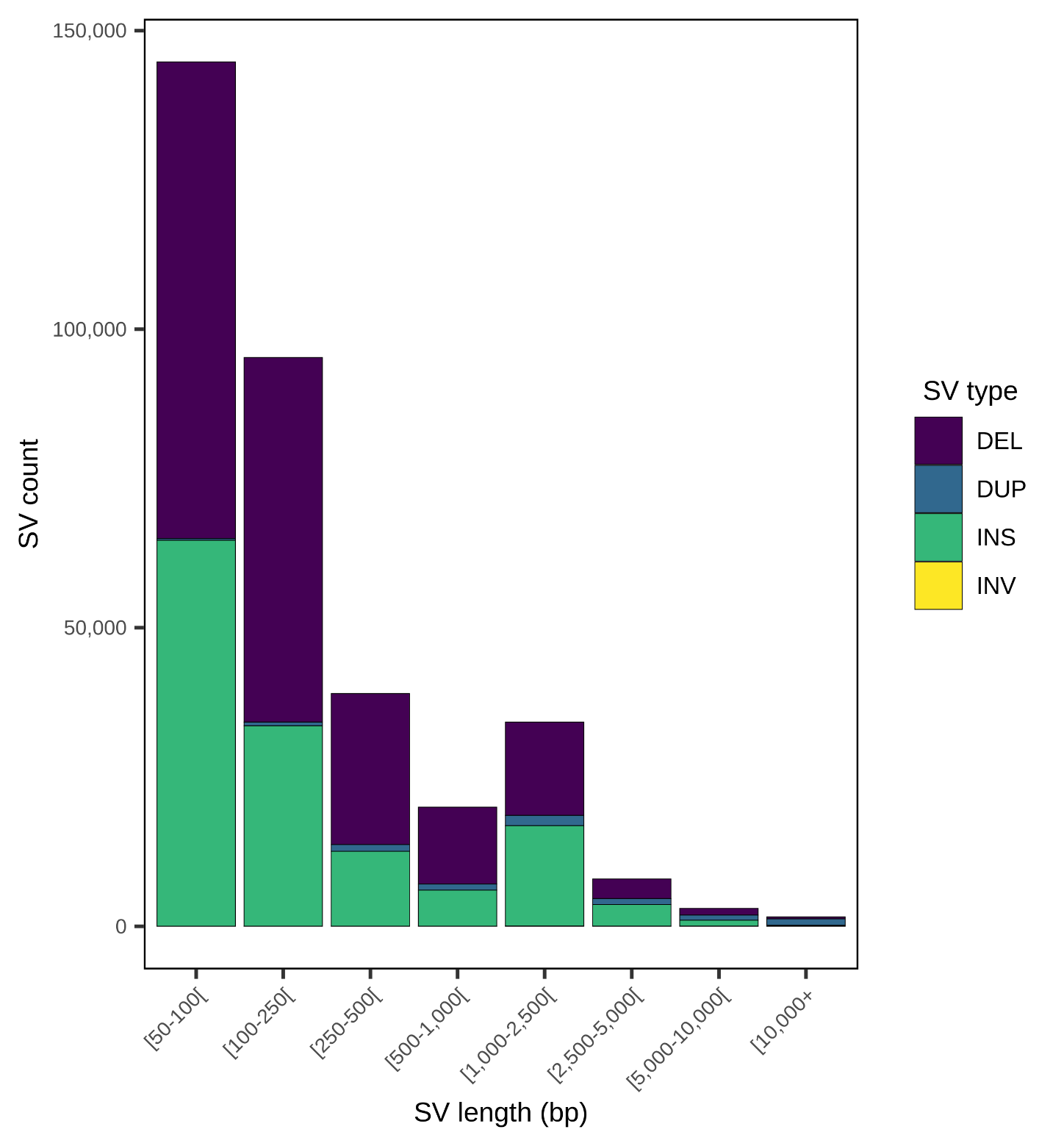

**Supplementary Figure 4.** Number of SVs in the filtered long-read SV set (LR SVs) by type and length, obtained by merging calls across the three long-read-based callers (Sniffles, SVIM and NanoVar) and filtering for a minimum of two supporting tools and a minimum length of 50 bp (DEL: deletions; DUP: duplications; INS: insertions; INV: inversions).

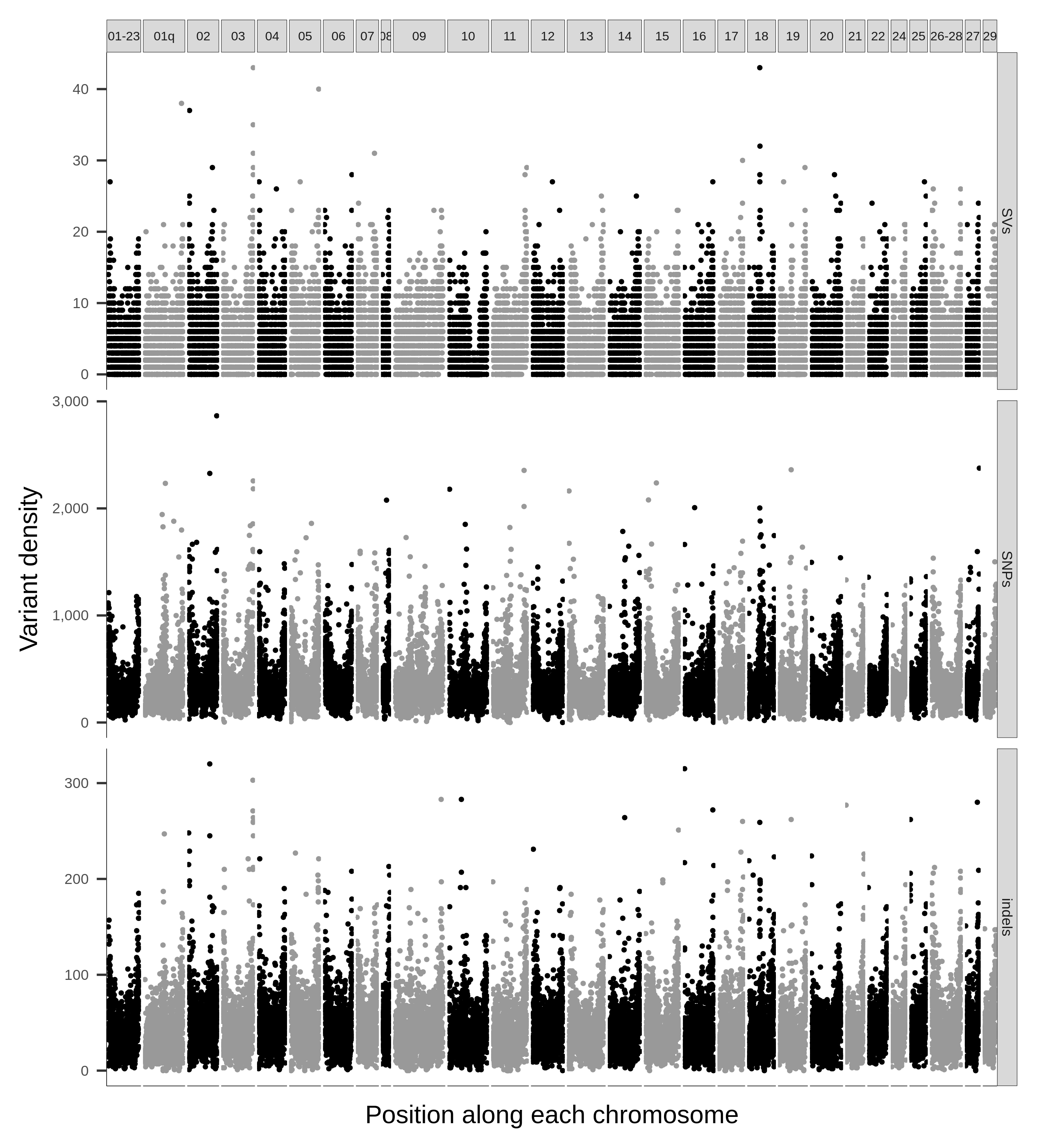

**Supplementary Figure 5.** Variant density, expressed in number of variants per 100-kb window along the genome for SVs (top), SNPs (middle) and small indels (bottom). Each vertical panel represents one chromosome.

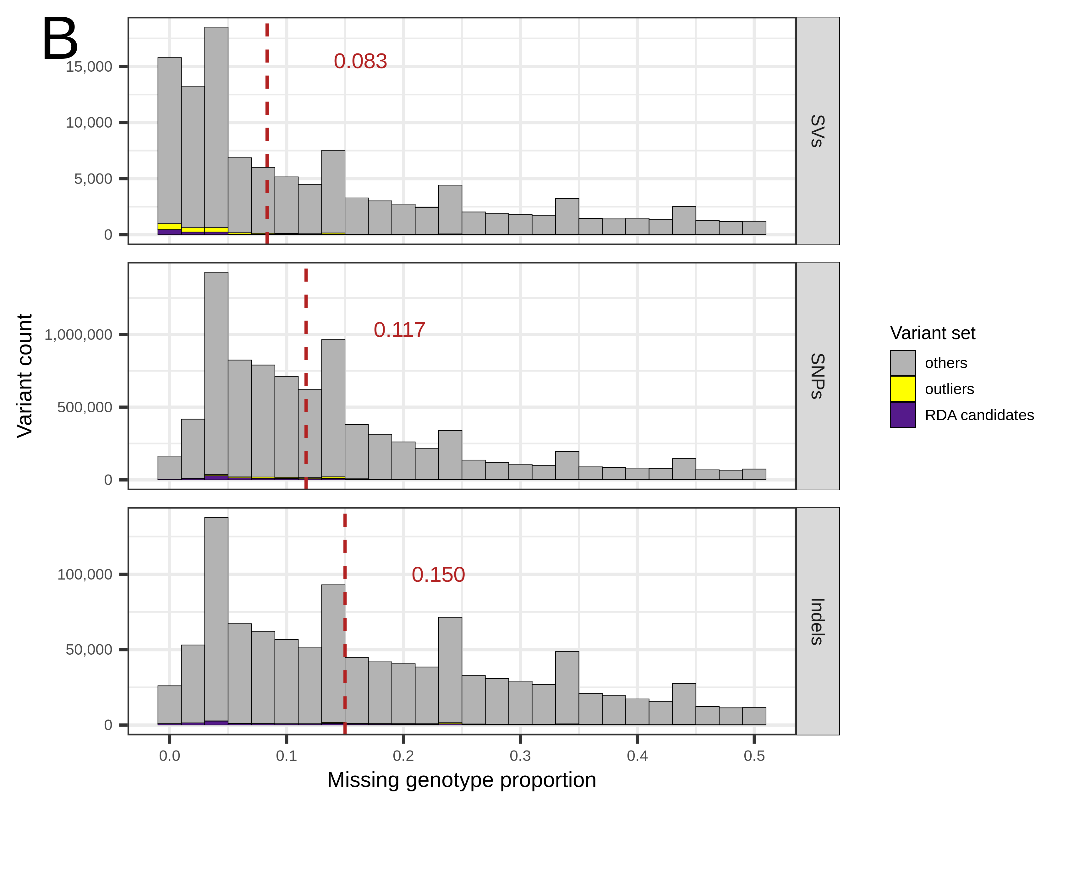

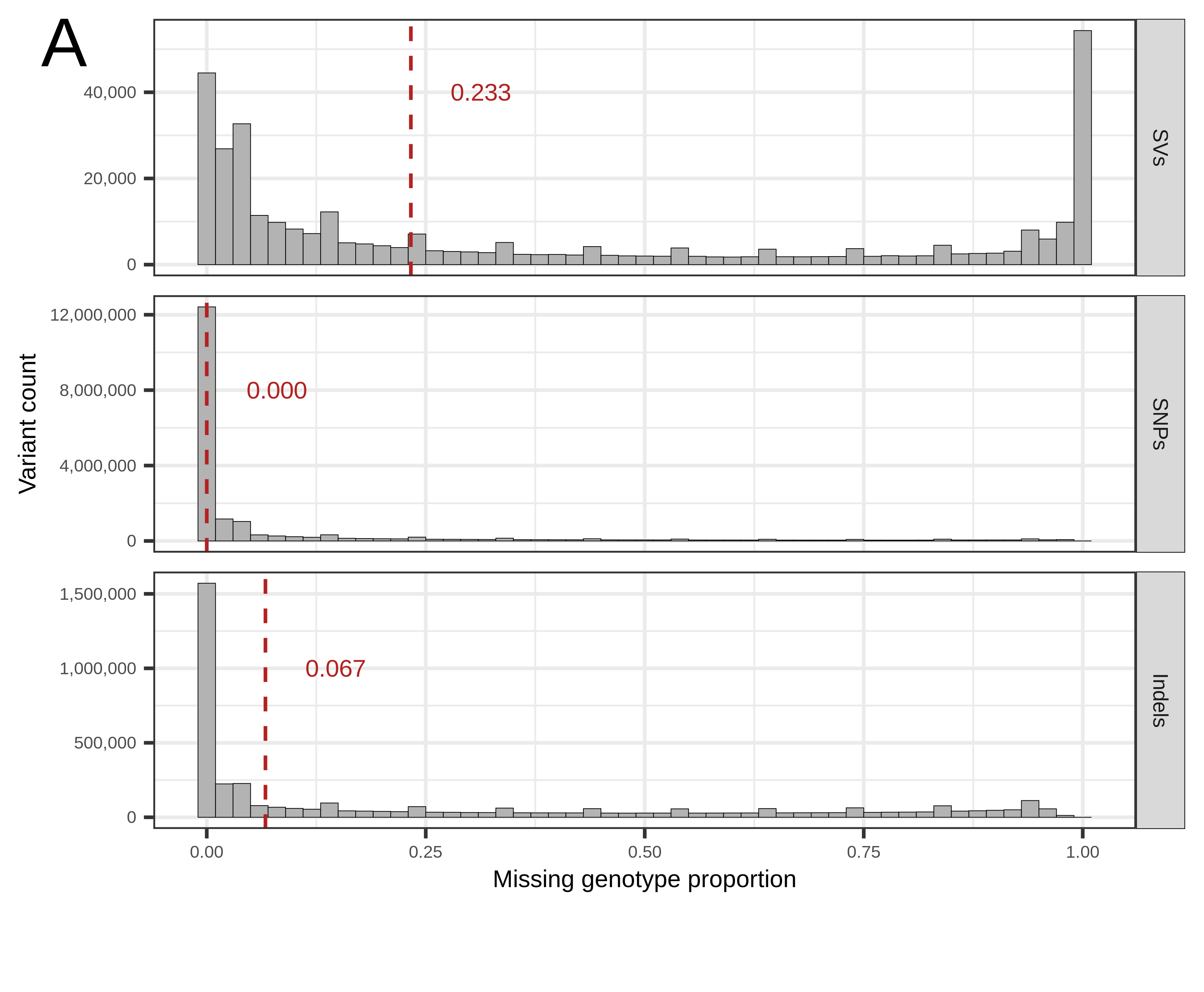

**Supplementary Figure 6.** Variant count by missing data proportion across SVs, SNPs and indels, **A)** for raw genotyped variants, and **B)** for filtered genotyped variants, colored by set, (outliers, RDA candidates and other variants, e.g., not outliers nor RDA candidates). The dotted red lines indicate the median missing data proportion. Two filters were used, one on the proportion of missing genotypes (F_MISS), which had to be lower than 50 %, and one on minor allele frequency (MAF) that had to be between 0.05 and 0.95. Both filters were applied together to produce the final filtered variant sets.

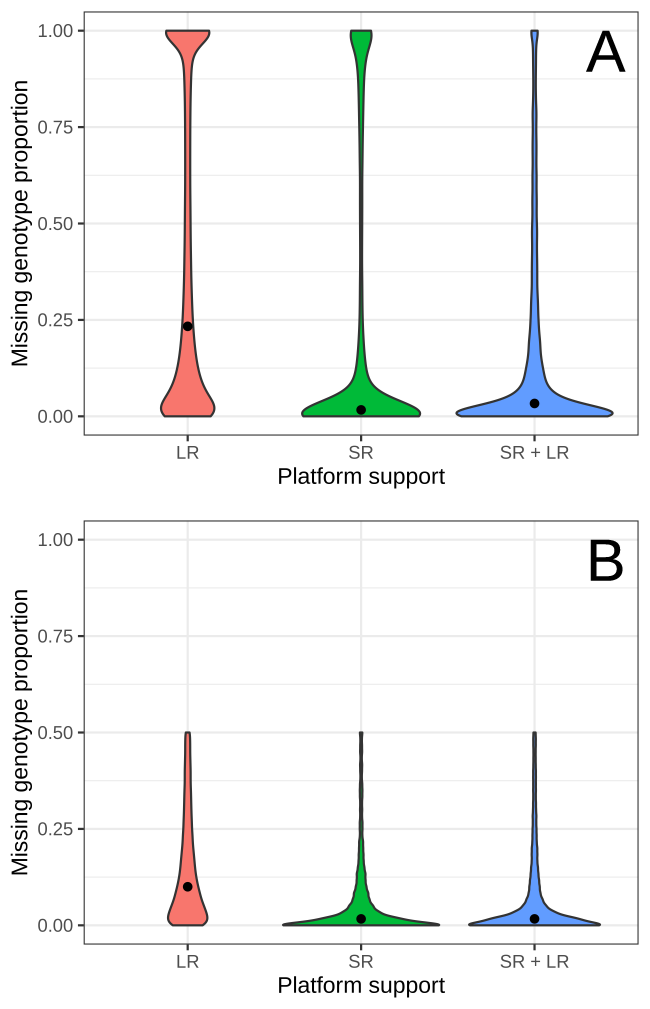

**Supplementary Figure 7.** Per-SV missing data proportion distribution by sequencing platform (SR: short-read; LR: long-read; SR + LR: short- and long-read) for **A)** raw genotyped SVs and **B)** filtered genotyped SVs. Sequencing platform information was inferred by matching genotyped SVs with known candidate SVs as described in Supplementary Methods 1.

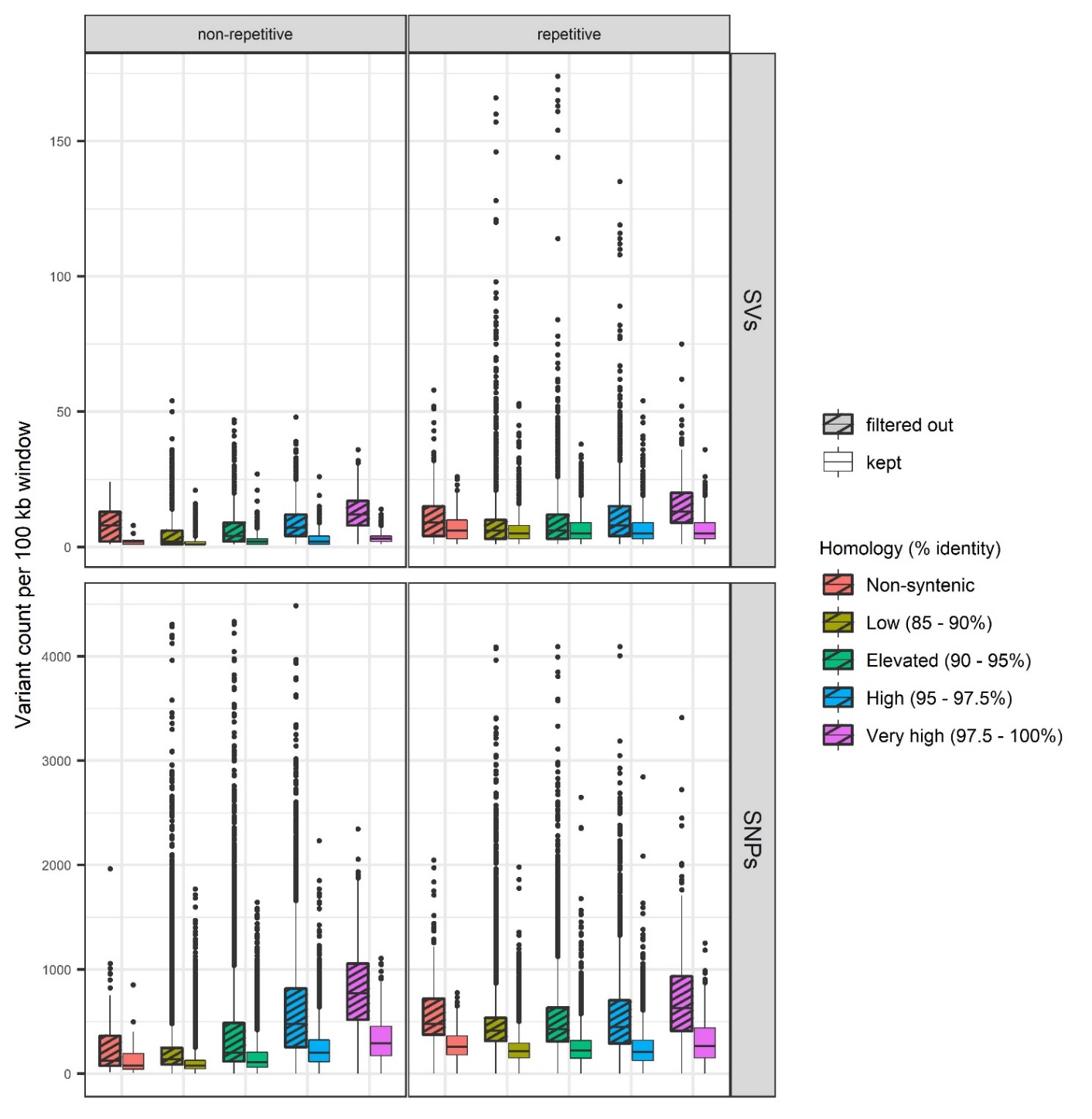

**Supplementary Figure 8.** Filtered and excluded genotyped SV (top) and SNP (bottom) count by 100-kb windows across different genomic features (repeated or non-repeated regions, duplicated/syntenic or non-duplicated regions). Syntenic regions are classified by their homology level, e.g., the percentage of identity between a given region and its duplicated counterpart/the corresponding duplicate region. Filtered out variants did not meet the requirements on genotype quality, depth, minor allele frequency and missing data proportion. Outliers (windows with more than 180 SVs or more than 4500 SNPs) were excluded for clearer visualization.

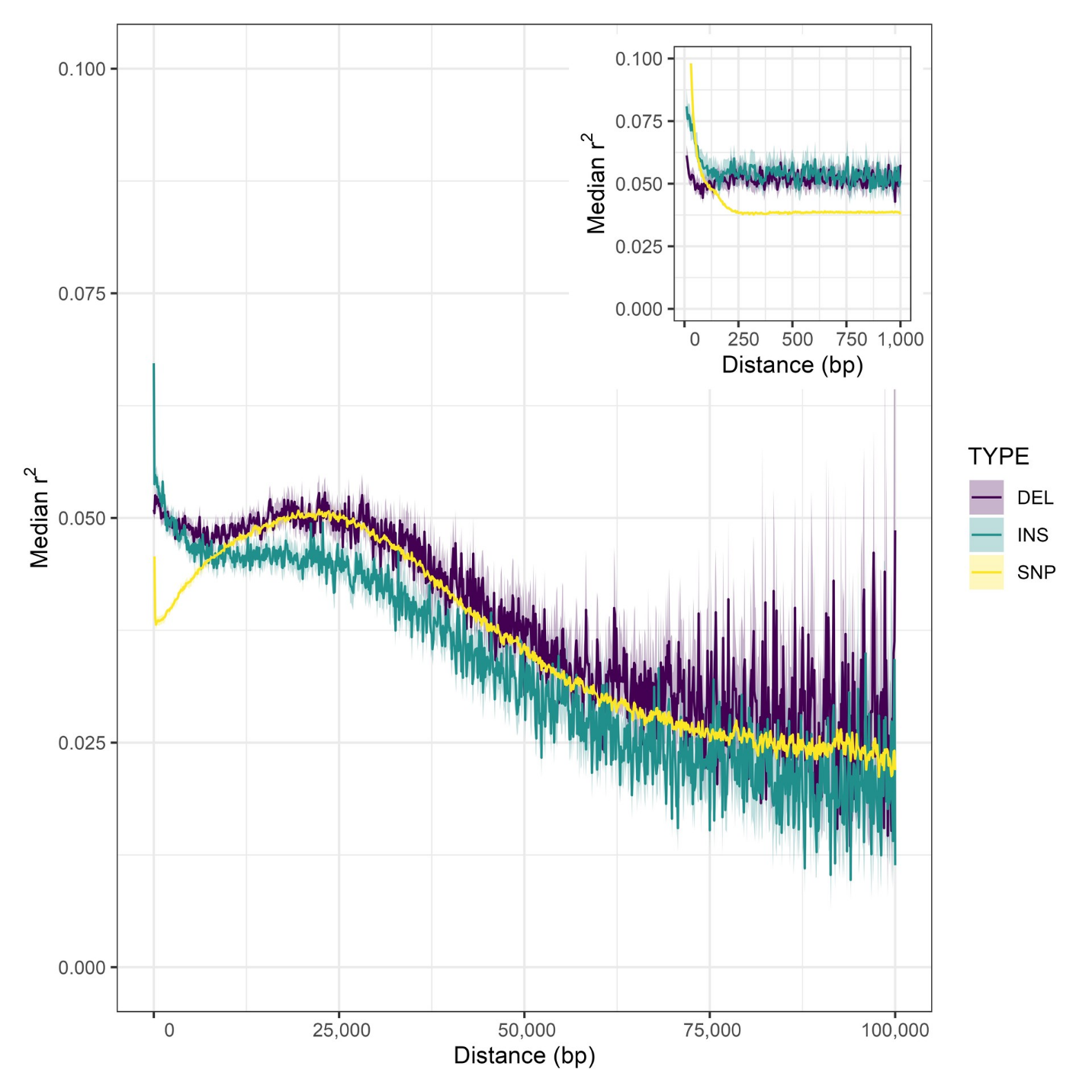

**Supplementary Figure 9.** Linkage disequilibrium (LD) decay between filtered SVs and SNPs (SV-SNP) and between SNPs (SNP-SNP). Median pairwise *r^2^* in 100bp distance bins is shown in the main panel. Inset panel represents a zoomed-in region on the first 1000 bp pairwise distance grouped over 5 bp bins. Shaded areas around lines represent the confidence interval inferred from bias-corrected bootstrap (*N* = 1,000) over a maximum of 150,000 pairs per distance bin. Inversions and duplications were excluded due to their insufficient number.

### Supplementary tables 1 to 4

**Supplementary Table 1.** Effect of various missing genotype proportion thresholds on remaining variant count after genotyping for each variant type. All other filtering criteria were equivalent to those used in the manuscript (e.g., biallelic sites with a minimal genotype quality of 5 and minimum depth of 4 in genotyped samples, and minor allele frequency between 0.05 and 0.95). The filtering threshold used (0.5) and the values reported in the manuscript are shown in bold characters.

|  | **SVs** | |  | **SNPs** | |  | **Indels** | |
| --- | --- | --- | --- | --- | --- | --- | --- | --- |
| **Maximum missing data proportion threshold** | **Remaining** | **Fraction of raw count** |  | **Remaining** | **Fraction of raw count** |  | **Remaining** | **Fraction of raw count** |
| None | 344,468 | 1 |  | 19,040,143 | 1 |  | 4,086,767 | 1 |
| 0.75 | 132,135 | 0.384 |  | 9,749,216 | 0.512 |  | 1,248,245 | 0.305 |
| 0.625 | 123,580 | 0.359 |  | 9,238,907 | 0.485 |  | 1,163,163 | 0.285 |
| **0.5** | **115,907** | **0.336** |  | **8,777,832** | **0.461** |  | **1,089,321** | **0.267** |
| 0.375 | 105,495 | 0.306 |  | 8,177,505 | 0.429 |  | 973,982 | 0.238 |
| 0.25 | 93,417 | 0.271 |  | 7,430,532 | 0.390 |  | 784,729 | 0.192 |
| 0.125 | 70,010 | 0.203 |  | 4,954,229 | 0.260 |  | 454,475 | 0.111 |

**Supplementary Table 2.** Merged SV count by type and sequencing platform, with corresponding proportions relative to the total 361,107 merged SVs (SR: short-read; LR: long-read; SR + LR: short and long-read). These represent putative SVs in the genomes of Romaine and Puyjalon salmon, prior to genotyping and filtering (DELs: deletions; DUPs: duplications; INSs: insertions; INVs: inversions).

| **SV type** | **Platform** | | | | | | | | **Per-type total** |
| --- | --- | --- | --- | --- | --- | --- | --- | --- | --- |
|  | **SR** | |  | **LR** | |  | **SR + LR** | |  |
|  | **Count** | **Proportion** |  | **Count** | **Proportion** |  | **Count** | **Proportion** |  |
| **DELs** | 13,972 | 0.065 |  | 178,690 | 0.837 |  | 20,789 | 0.097 | 213,451 |
| **DUPs** | 305 | 0.038 |  | 7,692 | 0.960 |  | 13 | 0.002 | 8,010 |
| **INSs** | 101 | 0.001 |  | 137,555 | 0.994 |  | 748 | 0.005 | 138,404 |
| **INVs** | 1,034 | 0.833 |  | 129 | 0.104 |  | 79 | 0.064 | 1,242 |

**Supplementary Table 3.** Summary of missing data statistics in raw and filtered variants sets. Raw SNPs and indels correspond to unfiltered variants called by the *SNPs_indels_SR* pipeline. Raw SVs correspond to the SVs that were genotyped through the *genotype_SVs_SRLR* pipeline, prior to filtration. Filtering was done on two levels: on a per-sample basis, e.g., individual genotypes that did not meet minimum allelic depth and genotype quality requirements were assigned the genotype "./.", and on the population level, e.g., on minor allele frequency and proportion of missing genotypes.

| **Variant type** | **Raw** | |  | **Filtered** | |
| --- | --- | --- | --- | --- | --- |
|  | **Count** | **Mean proportion of missing data per variant** |  | **Count** | **Mean proportion of missing data per variant** |
| **SVs** | 344,468 | 0.408 |  | 115,907 | 0.135 |
| **SNPs** | 19,040,143 | 0.092 |  | 8,777,832 | 0.144 |
| **Indels** | 4,086,767 | 0.260 |  | 1,089,321 | 0.183 |

**Supplementary Table 4.** Effect of population-level filters on variant count and proportion of remaining variants relative to the number of raw, unfiltered variants (None column). Two filters were used, one on the proportion of missing genotypes (F_MISS), which had to be lower than 50 %, and one on minor allele frequency (MAF) that had to be between 0.05 and 0.95. Both filters were applied together to produce the final filtered variant sets.

| **Variant type** | **Filters** | | | | | | | | | |
| --- | --- | --- | --- | --- | --- | --- | --- | --- | --- | --- |
|  | **None** |  | **F_MISS** | |  | **MAF** | |  | **F_MISS & MAF** | |
|  | **Count** |  | **Count** | **Proportion of raw count** |  | **Count** | **Proportion of raw count** |  | **Count** | **Proportion of raw count** |
| **SVs** | 344,468 |  | 214,519 | 0.623 |  | 155,007 | 0.450 |  | 115,907 | 0.336 |
| **SNPs** | 19,040,143 |  | 14,248,090 | 0.748 |  | 10,880,580 | 0.571 |  | 8,777,832 | 0.461 |
| **Indels** | 4,086,767 |  | 1,289,966 | 0.316 |  | 1,478,244 | 0.362 |  | 1,089,321 | 0.267 |

### Supplementary tables 5 to 10 legends

Supplementary tables 5 to 10 are provided in a separate Excel file.

**Supplementary Table 5.** Summary of the 108 significantly enriched GO terms for genes overlapping SV intersection outliers, clustered by REVIGO.

**Supplementary Table 6.** Summary of the 27 significantly enriched GO terms for genes overlapping SV RDA candidates, clustered by REVIGO.

**Supplementary Table 7.** Summary of the 223 significantly enriched GO terms for genes overlapping SNP intersection outliers, clustered by REVIGO.

**Supplementary Table 8.** Summary of the 528 significantly enriched GO terms for genes overlapping SNP RDA candidates, clustered by REVIGO.

**Supplementary Table 9.** Summary of the 212 significantly enriched GO terms for genes overlapping indel intersection outliers, clustered by REVIGO.

**Supplementary Table 10.** Summary of the 292 significantly enriched GO terms for genes overlapping indel RDA candidates, clustered by REVIGO.

### Supplementary tables 11 to 13

**Supplementary Table 11.** Assignment of the 115,907 genotyped and filtered SVs to a known putative SV, by SV type and sequencing platform (SR: short-reads; LR: long-reads; SR + LR: short and long-reads). 114,572 SVs were successfully matched to a known putative SVs based on position and alternate sequence length, as described in Supplementary Methods 1. The remaining 1,335 SVs could not be confidently assigned to a putative SV, either because they matched more than one putative SV or because no putative SV had equivalent position or alternate sequence length (DELs: deletions; DUPs: duplications; INSs: insertions; INVs: inversions).

| **Platform** | **Matched SVs** | | | | |  | **Unmatched or ambiguous** |
| --- | --- | --- | --- | --- | --- | --- | --- |
|  | **SV type** | | | | **Per-platform total** |  |  |
|  | **DELs** | **DUPs** | **INSs** | **INVs** |  |  |  |
| **SR** | 5,544 | 38 | 57 | 42 | 5,681 |  | 1,335 |
| **LR** | 45,297 | 352 | 48,750 | 11 | 94,410 |  |  |
| **SR + LR** | 13,795 | 0 | 678 | 8 | 14,481 |  |  |
| **Per-type total** | 64,636 | 390 | 49,485 | 61 | 114,572 |  |  |

**Supplementary Table 12.** Concordance between the candidate consensus genotype (i.e. called by SV callers) and the vg genotype across sequencing platforms (SR: short-reads; LR: long-reads) for genomes sequenced in both short and long reads, for raw genotyped and filtered (on MAF and F_MISS) genotyped SVs. Values represent the average proportion of concordant, non-concordant or unknown concordance across the four samples for each dataset. A consensus genotype was determined if the same genotype was called by at least two tools by platform, which was then compared to the genotype outputted by vg for the same SV in a given sample. If all three callers outputted different genotypes for a given SV in a given sample, then no consensus could be determined and the concordance with the vg genotype is unknown. Sequencing platform information was inferred by matching genotyped SVs with known candidate SVs as described in Supplementary Methods 1.

| **Genotype concordance** | **LR** | |  | **SR** | |
| --- | --- | --- | --- | --- | --- |
|  | **Raw** | **Filtered** |  | **Raw** | **Filtered** |
| **Concordant** | 0.376 | 0.325 |  | 0.670 | 0.790 |
| **Non-concordant** | 0.600 | 0.651 |  | 0.250 | 0.109 |
| **Unknown** | 0.024 | 0.024 |  | 0.079 | 0.100 |

**Supplementary Table 13.** Very large SVs (>30 kb) count by sequencing platform support (SR: short-read; LR: long-read; SR + LR: short- and long-read) across the SV candidates, the raw genotyped and the filtered genotyped SV sets. SV count by type is indicated in parentheses below platform support count in each cell (DELs: deletions; DUPs: duplications; INVs: inversions). Sequencing platform information was inferred by matching genotyped SVs with known candidate SVs as described in Supplementary Methods 1. All genotyped filtered SVs were deletions.

| **Dataset** | **Platform support** | | | **Total** |
| --- | --- | --- | --- | --- |
|  | **SR** | **LR** | **SR + LR** |  |
| **Candidates** | 437 (179 DELs; 21 DUPs; 237 INVs) | 351 (63 DELs; 268 DUPs; 20 INVs) | 16 (7 DELs; 1 DUP; 8 INVs) | 804 |
| **Genotyped, raw** | 96 (64 DELs; 13 DUPs; 19 INVs) | 282 (38 DELs; 243 DUPs; 1 INV) | 6 (4 DELs; 1 DUP; 1 INV) | 384 |
| **Genotyped, filtered** | 2 | 4 | 0 | 6 |

### Supplementary Methods 1. Inference of genotyped SVs properties by assignment to a known putative SV

We used pangenome-based genotyping to genotype SVs called from either long or short reads across all 60 samples using short-read data and the vg toolkit (Hickey et al., 2020). To do so, we built a variant-aware pangenome from the list of putative SVs, supplied as a VCF file. This VCF file was produced by the pipeline merge_SVs_SRLR (https://github.com/LaurieLecomte/merge_SVs_SRLR) and formatted at the initial preparation step of the genotyping pipeline, and included various information on each putative SV besides chromosome, position, reference and alternate allele sequences, such as SV type, length, end position and platform support (e.g., short and/or long reads).

However, vg does not carry on these optional INFO fields throughout the genotype calling procedure, as it is performed by reference to the variant-aware graph structure, and not to the putative SVs per se. vg therefore outputs a new ID and minimal information for each genotype call, e.g., chromosome, position, reference and alternate alleles sequence fields.

In order to estimate the amount of genome base pairs covered by SVs (especially insertions) and the proportion of successfully genotyped SVs that were originally called from long and/or short reads, we attempted to infer info on SV length and platform support by matching genotyped SVs with known putative SVs, based on position and reference and alternate alleles sequence size. This procedure is performed using the script compare_summarize_plot.sh (from the genotype_SVs_SRLR pipeline; https://github.com/LaurieLecomte/genotype_SVs_SRLR), briefly explained below, along with the putative SVs VCF file and the filtered genotyped SVs VCF file.

First, for each putative inversion, we retrieved SVLEN and END fields from the short-read SV (from the SVs_short_reads pipeline; https://github.com/LaurieLecomte/SVs_short_reads) and the long-read SV (from the SVs_long_reads pipeline; https://github.com/LaurieLecomte/SVs_long_reads) sets, because these were lost when merging short and long-read SV sets with Jasmine (add_missing_INV_info.R).

Second, in R, we computed reference and alternate allele length for all putative and genotyped SVs. We then assigned a known putative SV to genotyped SV by performing a conditional merge of both datasets on chromosome, position, as well as on reference and alternate allele sequence lengths, allowing a 5 bp window (infer_info_genotyped.R).

114,572 genotyped SVs could thus be confidently matched to a single known putative SVs (Suppl. Table 11), from which retrieved SV type, SV length and platform support. In addition, 1335 genotyped SVs either could not be matched to any putative SVs, or had multiple matches and were therefore deemed ambiguous.

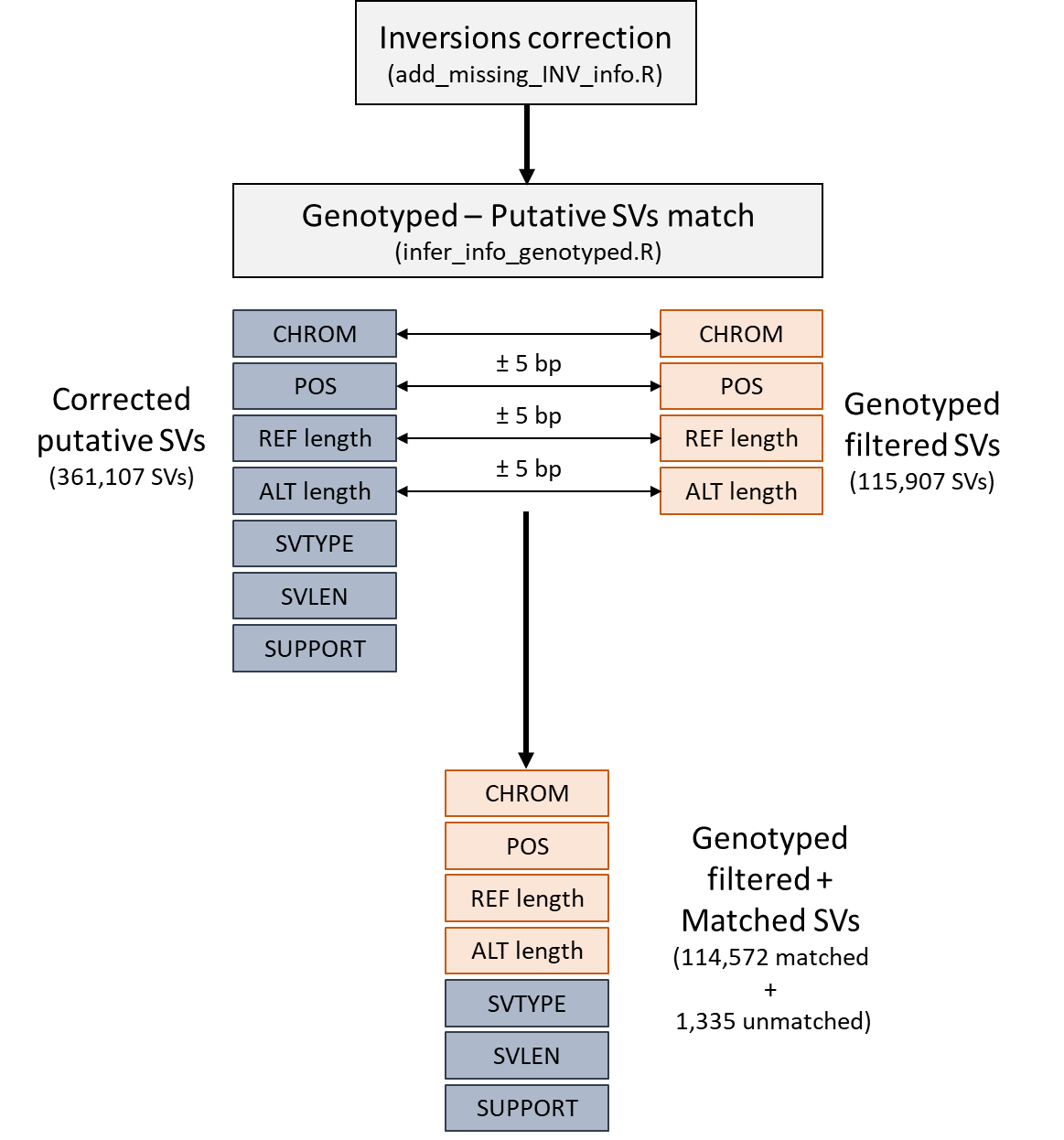
